# The lattice scales of grid-cell activity

**DOI:** 10.64898/2026.08.07.743499

**Authors:** Arseniy Veselov, Anian Kerscher, Martin Stemmler, Andreas V.M. Herz

**Author notes:** equal contribution (Arseniy Veselov, Anian Kerscher).

## Abstract

Grid cells in the rodent medial entorhinal cortex (MEC) form a multi-scale representation of the animal’s environment and are thought to be a signature of continuous-attractor networks.Experiments suggest that grid scales are discrete and arranged in a geometric progression with constant scale ratios, as predicted by efficient-coding theories that posit that the grid-cell system is optimized for spatial resolution. This normative explanation has been challenged by a developmental theory whereby grid-cell modularity arises from pattern formation in a system with smooth parameter gradients. As a consequence, scale ratios should *not* be constant, but vary by module. However, as we show here, the experimental data chosen to support this mechanistic theory fail to do so. Publicly available large-scale grid-cell datasets could provide a clear benchmark and further our understanding of the neuronal basis of spatial navigation. We, therefore, reanalyzed such recordings and found that they refute the developmental theory. Instead, the measured scale ratios agree with the geometric-progression hypothesis and set strict bounds for any future grid-cell theory.

---

As a function of animal location, grid cells exhibit multiple firing fields that form a hexagonal lattice across the explored environment.^1^ The grid cells of an animal are organized into a small number *n* of modules *M*_*m*_, 1 ≤ *m* ≤ *n* (Fig. 1a).^2–5^ Within a module, the firing fields of all grid cells share a common grid scale *λ*_*m*_ and aligned lattice orientations but vary in their spatial phase. In rats, measured grid scales range from 30-40 cm in dorsal MEC to about 180 cm in ventral MEC.^1–5^ As we will show, the experimental grid-scale values and their ratios across neighboring modules (Fig. 1b) set critical constraints for theories on the development and function of the grid-cell system. For concreteness, modules will be ordered by decreasing scale so that *m* = 1 labels the largest-scale module and *λ*_*m*_ *> λ*_*m*+1_.

**Figure 1.**
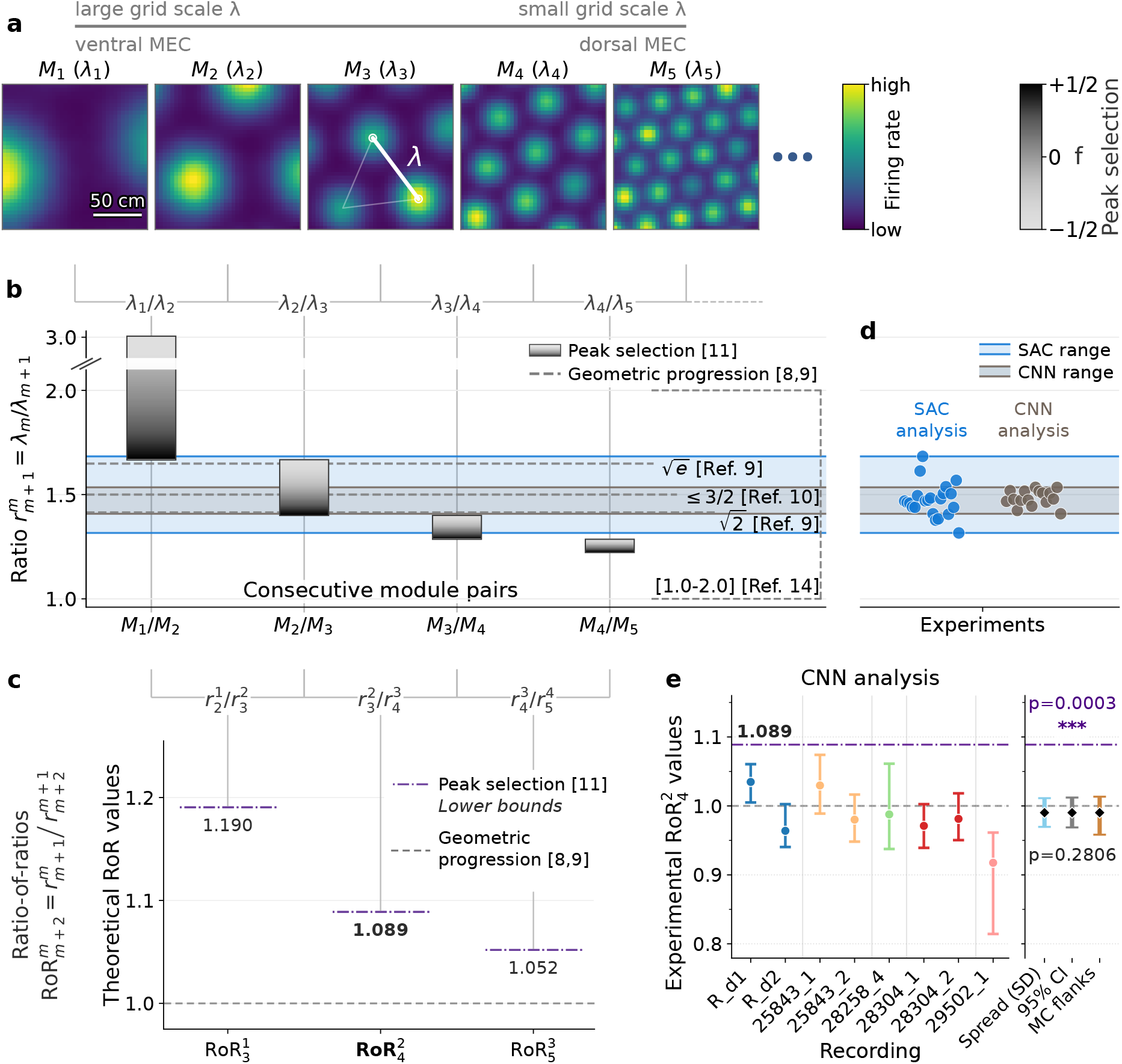
Predicted and measured grid-scale ratios and ratios-of-ratios. **a**, Idealized firing-rate maps for five consecutive modules with grid scales *λ*_1_ −*λ*_5_. **b**, Theoretical scale ratios for the four pairs of consecutive modules. Gray-scale bars: parameter-(f) dependent predictions from peak-selection theory.^11^ Dashed lines depict predictions from geometric-progression theory^9^ and population-vector decoding,^10^ and the vertical bracket indicates the geometric-attractor^14^ range. **c**, Conservative (f=1/2) lower peak-selection bounds for the three consecutive ratios-of-ratios. Dark dashed line: the geometric-progression prediction (RoR ≡ 1). **d**, Estimates from recordings with three modules. Dots show raw estimates, shaded bands are the observed ranges for spatial-autocorrelogram (SAC) and convolutional-neural-network (CNN) analyses. **e**, CNN estimates of 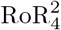 for eight recordings. Points and whiskers: raw estimates and central bootstrap 95% intervals, colored by animal. Rat R was recorded by Gardner et al.,^12^ the four other animals by Vollan et al.^13^ Reference lines indicate RoR=1 and the peak-selection bound 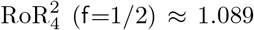. In the population summary, black diamonds mark the reliability-weighted geometric mean; “Spread” shows weighted SD of the animal estimates around the population mean, “95% CI” is the mean’s HC1 *t* confidence interval, and “MC flanks” depicts the median lower and upper animal-level Monte Carlo uncertainty flanks. The RoR population estimate is 0.99 (95% CI, 0.97–1.01), consistent with geometric-progression theories but disproving the explanation based on peak selection.

Because grid-cell activity is periodic, the cells’ spatial representation remains ambiguous at the single-module level. Combining multiple modules with different grid scales overcomes this limitation^6^ and yields highly efficient population codes^6–9^ that outperform place-cell-like representations in terms of spatial range^6, 7^ and resolution.^8, 9^ Optimal resolution is achieved when the scale ratio 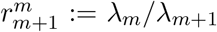 between adjacent modules *M*_*m*_ and *M*_*m*+1_ is constant, 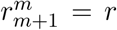, across modules,^8, 9^ so that grid scales follow a geometric progression. Population-vector decoding predicts that *r* should not exceed 3*/*2 to avoid catastrophic decoding errors,^10^ whereas minimizing neuron numbers for fixed spatial resolution gives 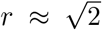 or 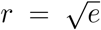, depending on the decoder.^9^

These predictions are to be compared with the developmental theory by Khona et al.^11^. The authors propose a morphogenetic process that combines pattern formation and gradient-based positional mechanisms, creating distinct modules with discrete grid scales, as do the geometric-progression theories.^8, 9^ However, according to the “principle of peak-selection” put forward by Khona et al., scale ratios should *not* be constant^8, 9^ but decrease with increasing module index.^11^ Specifically, the ratios should obey

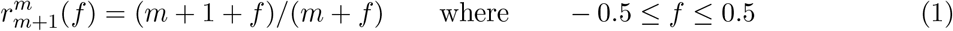

is a free parameter that depends on the synaptic connectivity and grid-cell time constants.

For *f* = 0, equation (1) simplifies to 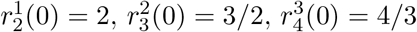, etc. Indeed, in the data of Stensola et al.^3^ showcased by Khona et al., there is one animal (Rat 14257) whose scale ratios (1.514, 1.343, 1.247) deviate from the ideal values on average by less than 0.7% (RMSE) if one starts with *M*_2_ as the largest-scale module (presuming that no neurons from module *M*_1_ were recorded). For two other animals (Rats 14147 and 15708), the authors assume that the largest-scale module is *M*_1_, without explanation. Optimizing the parameter *f*, they obtain scale ratios that differ from the experimental values on average by 17.8% and 17.9%, respectively (RMSE). Alternatively, within the geometric-progression framework, the ratios should scatter around some animal-specific mean. Measured against this hypothesis, the scale ratios of the three animals show average mismatches of 8.1%, 5.6%, and 6.0% (RMSE). We conclude that even for the data selected by Khona et al., fits from geometric-progression theories are almost twice as good (average RMSE: 6.6% vs 12.1%) as those from the peak-selection theory.

To go beyond such particular observations, we wondered about the general implications of equation (1). Independent of the choice for *f*, scale ratios should decrease with increasing *m* (Fig. 1b), in stark contrast to the constant ratio *r* predicted from optimizing spatial resolution. Taking *f* into account, equation (1) implies that successive scale ratios should lie in bands that shift downwards, 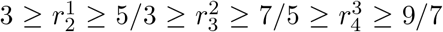, etc. The ratio of ratios, 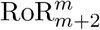, defined as 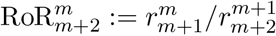, takes its smallest value for *f* = 1*/*2, resulting in the following bounds: 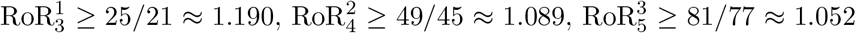, etc (Fig. 1c). Comparing these predictions with those from geometric progression, for which 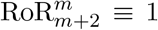, provides a decisive test to distinguish the two mutually exclusive hypotheses.

Precise grid-scale measurements are essential for this test. We therefore re-analyzed Neuropixel data with large spike counts per grid cell, recorded by Gardner et al.^12^ and Vollan et al.^13^ (Supplementary Fig. 1), and used the grid-cell activity’s spatial autocorrelation (SAC) to calculate lattice scales (Supplementary Information sections S4, S5). For these datasets, 21 of 22 scale ratios 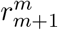 are below 5*/*3 (Fig. 1d, Supplementary Table 4). According to the peak-selection theory, therefore, none of those grid cells belongs to an *M*_1_ module.

We then examined grid-scale ratios from recordings with three adjacent grid modules, i.e., two successive scale ratios, as needed for the RoR calculation. The larger of the two ratios fell between 7*/*5 and 5*/*3, with only two exceptions (Fig. 1d), so, following Khona et al., we labeled the modules as *M*_2_, *M*_3_, and *M*_4_, respectively.

From equation (1), 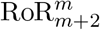 is closest to unity (and smallest) when *f* = 1*/*2. Indeed, 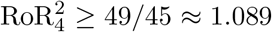 (Fig. 1c) must hold in the theory of Khona et al., yet the experimental 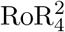 values and their bootstrap intervals remain below this bound in 9 of 11 recordings (Supplementary Fig. 5d); the two exceptions are the recordings with the fewest cells (Supplementary Discussion), whose wide bootstrap intervals overlap the bound. But the 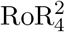 values are close to unity: the weighted population mean is 0.963 (SD 0.903–1.028; 95% CI 0.914–1.015), consistent with geometric progression (*p* = 0.136; HC1 *t*-test), yet clearly below the conservative peak-selection bound (*p* = 8.9 *×* 10^−4^; HC1 *t*-test).

For large grid scales (Fig. 1a, panels *M*_1_ and *M*_2_), the SAC analysis systematically under-estimates their size when too few firing fields fall completely within the arena explored by the animal, as confirmed by synthetic data (Supplementary Fig. 2a). This, in turn, can cause an underestimation of 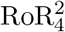 as shown by one example in Supplementary Fig. 5b.

We therefore repeated the entire analysis, replacing the SAC method by a convolutional neural network (CNN) trained to predict grid scales (Supplementary Information section S4). As expected, this yielded higher 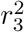 values (Supplementary Fig. 4) so that the ratios became virtually identical: 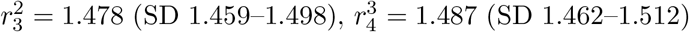. These values are consistent with predictions by Stemmler et al.^10^ (*r* ≤ 3*/*2) and mouse data (*r* = 1.49*±*0.08) from Kerekes et al.^5^, and close to a prediction by Wei et al.^9^ 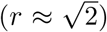 and the original observation (*r* = 1.42 *±* 0.02) of Stensola et al.^3^. The slightly larger values of Barry et al.^2^ (≈ 1.63) and Krupic et al.^4^ (≈ 1.56) are possibly due to limitations in sample size.

The CNN-estimate of 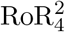 is 0.9902 (SD 0.9696–1.0112; Fig. 1e), consistent with RoR ≡ 1 from geometric progression^8, 9^ (*p* = 0.281; HC1 *t*-test) but clearly below the bound 1.089 from peak-selection theory (*p* = 2.7 *×* 10^−4^; HC1 *t*-test).

Our analysis refutes the peak-selection theory of Khona et al.^11^ as a viable account of how and why the rodent grid-cell system is organized in discrete modules. This does not question the mathematical beauty of this theory nor does it rule out alternative developmental theories such as that proposed by Kang and Balasubramanian,^14^ which predicts a constant scale ratio *r* between one and two, depending on model details. Our findings underscore that the geometric-progression hypothesis^8, 9^ offers the most parsimonious explanation for the experimentally observed modular structure and its implications for the impressive computational function^15^ of grid cells. The quantitative results impose tight constraints for any future grid-cell theory.

## Acknowledgements

We are grateful to Caswell Barry and Julija Krupic for sharing valuable data and thank Sarthak Chandra, Ila Fiete, Alexander Mathis and Edvard Moser for helpful comments on the manuscript.

