## Supplementary Material for "The lattice scales of grid-cell activity"

Supplementary Information

Arseniy Veselov*, Anian Kerscher*, Martin Stemmler, Andreas V.M. Herz

August 7, 2026

Grid-cell modules are organized according to the length scale of their grid fields along the dorso-ventral axis of medial entorhinal cortex. For this ladder of scales, starting from the largest to the smallest, theories predict either a constant scale ratio[8,9] or a decreasing scale ratio.[11] Experimental verification, the subject of this supplement, requires careful consideration: when the grid scales are large compared to the size of the arena, only a few grid fields are observed, some of which are partially occluded by the arena borders. Such grid scales are easily underestimated, which in turn will affect the estimated scale ratios. Here we describe the full analysis pipeline on both synthetic and real data to distinguish the two theories.

### Supplementary Methods

#### S1 Data sources, scope, and recording inventory

The empirical analyses focused on open-field Neuropixels recordings from Gardner et al.[12] and Vollan et al.[13]. These datasets contain spike trains from large, simultaneously recorded populations from medial entorhinal cortex, parasubiculum, and adjacent parahippocampal tissue, together with data from tracking the animal's two-dimensional position. Recording duration, spike count, and the number of retained cells vary across recordings and are summarized in Supplementary Fig. 1a,b. Here, "recording" denotes the open-field dataset associated with a single identifier from the experimentalists; separate recordings could occur on the same or on different days. Acquisition, probe placement, behavioral protocols, spike sorting, and animal procedures are described in the source publications and are not repeated here except where they affect the present analysis. We considered all open-field recordings in the two source datasets and restricted the analysis to cells labeled as grid cells in the respective sources. Source recording identifiers were preserved. For the Vollan et al. recordings, the numeric prefix identifies the animal and the suffix denotes a distinct recording; similarly, the recordings for the rat `R` acquired by Gardner et al. are denoted `R_d1` and `R_d2`. Repeated recordings from the same animal were retained as separate recording-level observations but were nested within each animal in the primary population analysis (Section S6).

The key advantage of these data for the present analysis is that many single recordings contain enough co-recorded grid cells to resolve the centers of three modules simultaneously. Earlier module sequences, including the data used in Khona et al.[11], were inferred from smaller samples and often required information to be combined across sessions, environments, or electrode positions[3]. Detecting the large-scale modules nevertheless remains difficult in a finite arena because only a small number of fields may be visible and the probe need not span the complete dorsoventral axis (Section S4).

**Conditional module nomenclature.** The present analysis identifies a triple of putative grid-scale module centers, ordered from largest to smallest, but does not establish their absolute anatomical indices. For comparison with the peak-selection theory proposed by Khona, Chandra, and Fiete[11], following the indexing used in the main text, this triple is written $(M_2, M_3, M_4)$. If the peak-selection theory does not describe the data, the labels carry no information about how many larger modules may exist. The normative geometric-progression theories considered here[8,9] likewise do not, by themselves, determine the absolute anatomical rank of an observed triple.

**Arena geometry.** Most recordings were obtained in a standard $150\,\text{cm} \times 150\,\text{cm}$ square open field. In `26018_2`, the animal foraged over an approximately circular region about 1.5 m in diameter. The recording was retained for the spatial autocorrelogram (SAC) ratio analyses (Section S5) but excluded from the convolutional neural network (CNN) ratio analysis because the CNN was trained using the $150\,\text{cm} \times 150\,\text{cm}$ arena geometry (Section S4).

**Recording eligibility.** After the cell-level grid-cell and estimator-specific quality-control steps described in Sections S2 and S4, a recording entered the ratio-of-ratios (RoR) analysis when Stage 2 (Section S5) recovered at least three module centers, subject to the CNN-specific arena-geometry exclusion described above. Eligibility was evaluated separately for the SAC and CNN analyses and was defined independently of the resulting ratios or their relation to either theoretical prediction. No recording yielded more than three fitted modules.

The final cohorts contain 11 recordings from 8 animals for SAC and 8 recordings from 5 animals for CNN (all five animals in the CNN cohort were also present in the SAC cohort). We did not impose a minimum number of cells assigned to each fitted module because no independent biological or benchmark-derived threshold justified a specific cutoff. Instead, instability under cell resampling is quantified by the recording bootstrap (Sections S6 and S7), and the resulting conditional uncertainty is incorporated into the reliability-weighted animal and population summaries (Section S6). Recordings with limited fitted-module support therefore remain visible and auditable rather than being removed post hoc. Supplementary Table 4 lists the included recordings, ordered module centers, adjacent ratios, and raw RoR values; Supplementary Table 5 reports the associated bootstrap distributions and retained-replicate counts.

**Analysis overview.** For each recording, we first constructed occupancy-corrected firing-rate maps and retained grid cells that passed the common grid-score criterion. Grid scale was then estimated independently with the SAC and CNN methods, using the corresponding estimator-specific quality-control procedure. The retained cell-scale estimates were analyzed in log space, candidate Gaussian mixture models were fitted, and the selected components were ordered from largest to smallest scale. Recordings in which three ordered components were recovered contributed the two adjacent scale ratios and their ratio of ratios. Recording-level uncertainty was quantified by resampling cells and refitting the complete module model; repeated recordings were subsequently combined for the same animal, and the resulting animal estimates were combined at the population level using regularized reliability weights. Cellwise simulations evaluated scale estimation, whereas separate three-module simulations tested whether the complete pipeline could recover known constant- and decreasing-ratio structures.

#### S2 Rate-map construction and basic grid-cell inclusion

Spike times were aligned to the animal's tracked position by temporal interpolation of the recorded trajectory and converted into occupancy-corrected two-dimensional firing-rate maps. The arena was represented as the fixed square domain $\mathcal{A} = [-0.75, 0.75] \times [-0.75, 0.75]$ m, centered at the origin. The spatial domain was discretized into a fixed $100 \times 100$ grid based on the absolute coordinates of the arena boundaries (identical across all datasets and recordings, including the circular arena). Position samples were binned into this predefined grid, and spike counts were accumulated in the same spatial bins. $S(x, y)$ denotes the binned spike-count map, $O(x, y)$ the corresponding occupancy map, and $M(x, y) \in \{0, 1\}$ the binary validity mask indicating all spatial bins within the physical boundaries of the arena.

Smoothing was performed in a boundary-aware manner. The spatial bandwidths for smoothing the spike and occupancy maps were set to be equal ($\sigma_s = \sigma_o$). A standard deviation of 4 cm was used, which, given a spatial bin size of 1.5 cm, corresponds to a discrete Gaussian kernel with $\sigma \approx 2.67$ bins. Each smoothed image was normalized by the effective kernel mass supported inside the valid arena:

$$\widetilde{S}_{\sigma_s}(x, y) = \frac{(G_{\sigma_s} * (SM))(x, y)}{(G_{\sigma_s} * M)(x, y)}, \tag{1}$$

and

$$\widetilde{O}_{\sigma_o}(x,y) = \frac{(G_{\sigma_o} * (OM))(x,y)}{(G_{\sigma_o} * M)(x,y)}. \tag{2}$$

This correction prevents ordinary convolution from attenuating near-boundary values simply because part of the Gaussian kernel lies outside the arena. It is important here, as without it the lattice at arena boundaries would be by design shifted inwards. The convolutions were performed in the spatial domain, using zero-padding for regions outside the defined arena mask to explicitly manage boundary effects.

The occupancy-corrected spatial rate map was then computed as

$$r(x,y) = \begin{cases} \frac{\widetilde{S}_{\sigma_s}(x,y)}{\widetilde{O}_{\sigma_o}(x,y)}, & \text{if } \widetilde{O}_{\sigma_o}(x,y) > O_{\min} \\ \text{NaN}, & \text{otherwise,} \end{cases} \tag{3}$$

where $O_{\min}$ is an occupancy floor used to exclude poorly sampled bins. It was defined as the 1st percentile of the smoothed occupancy, clamped between 0.01 and 0.1 seconds.

The same underlying rate-map representation was used for the spatial-autocorrelogram (SAC), convolutional neural network (CNN), and optional geometric estimators. This is important because any difference between the SAC and CNN estimates should then be attributable to the estimator, not to different preprocessing choices.

Grid-cell inclusion at this stage followed the cell labels and quality-control logic of the source datasets. For both the SAC and CNN methods, the initial cell selection from the complete grid-cell population was based on the grid score, calculated by applying the expanding annulus method[16,17] to the spatial autocorrelogram of the firing-rate map. The grid score was evaluated iteratively across a series of concentric annuli. The inner radius of the annular region was fixed at the edge of the central peak plus a spatial margin (9 cm, corresponding to 6 bins of size 1.5 cm). From this starting point, the outer radius expanded sequentially in steps of 1 bin up to a maximum boundary (the arena size minus a 9 cm margin). To determine the edge of the central peak, we computed the radial profile of the SAC. Specifically, for each concentric 1-bin-wide ring extending from the center, we calculated the mean autocorrelation value of all spatial bins within that ring. The central peak radius was then defined as the shortest distance from the center at which this mean autocorrelation either reached its first local minimum or dropped below a threshold of 0.2.

To evaluate rotational symmetry, the entire SAC was first rotated by 30°, 60°, 90°, 120°, and 150°. Then, for each defined annulus, the Pearson correlation coefficient was calculated between the unrotated and each rotated SAC. This computation was restricted strictly to the spatial bins falling within the specific annular region that contained valid, non-missing data in both arrays. For a given outer radius, the intermediate grid score was computed as the minimum correlation at the symmetry angles (60° and 120°) minus the maximum correlation at the offset angles (30°, 90°, and 150°). The final grid score assigned to the cell was the maximum value obtained across all evaluated annuli. For subsequent analyses, only grid cells with a final grid score of $\geq 0.2$ were retained.

To pass the SAC quality-control criteria, a cell was required to exhibit at least 6 distinct autocorrelation peaks, excluding the central peak. Following the procedure described by Stensola et al.[3], these six innermost peaks were identified to define the three grid axes. Specifically, the axes were defined by the vectors from the SAC center to the first six local maxima surrounding the central peak. Based on these axes, two geometric filters were applied to ensure grid regularity: (1) the inter-axis angles between all three adjacent grid axes had to be strictly between 30° and 90°, and (2) all pairwise ratios of the three grid-axis lengths had to fall between 0.5 and 2.

For the CNN estimator, aside from the initial grid-score threshold, no additional geometric quality-control criteria were applied. We deliberately omitted further filters because the probabilistic nature of the CNN inherently quantifies the uncertainty of its scale and orientation predictions.

### S3 Synthetic data generation and benchmark rationale

Synthetic data served two distinct validation purposes and were not intended as a mechanistic grid-cell model. First, a large cellwise simulation bank was used to train the CNN and to evaluate, on held-out stochastic realizations, whether the CNN and SAC estimators recovered known lattice spacing over the finite-arena range relevant to the experimental recordings (Section S4.2). Second, separate three-module populations were used to test whether the complete downstream pipeline – cellwise estimation, logarithmic GMM fitting, adjacent-ratio construction, and ratio-of-ratios calculation – could distinguish a locally constant-ratio sequence from a peak-selection-like unequal-ratio sequence. The three-module populations were not used to train the CNN or to select its architecture, quality-control criteria, GMM settings, or statistical summaries.

#### S3.1 Common generative and observation model

This subsection defines the lattice generator, empirical trajectory, and Poisson observation process shared by the two synthetic benchmarks.

Each synthetic cell was generated from a hexagonal lattice with prescribed spacing $\lambda$, orientation $\theta$, and spatial phase. Lattice fields were represented by isotropic Gaussian bumps on a $201 \times 201$ grid within a nominal $150\,\text{cm} \times 150\,\text{cm}$ arena. The lattice was generated within a 40 cm surrounding margin, allowing field centers outside the nominal arena to remain part of the lattice. Together, this formed the spatial template $\omega_i$ for cell $i$.

All simulations used an empirical open-field trajectory from Gardner rat `R`, day 1 ($\sim$ 2 h 21 min). This long recording provided dense sampling of the standard arena and was used to impose realistic occupancy inhomogeneity and finite spatial coverage. The same recording also supplied the empirical spike-count banks used in both benchmarks and, for the three-module benchmark, the module-specific banks of retained-cell spatial phases. Thus, the benchmarks shared the trajectory, arena geometry, and observation model, while their cell-specific parameters were generated according to the benchmark-specific rules given below. The simulations therefore reproduce selected sampling conditions of the recordings while retaining known ground-truth lattice spacings.

Spikes were generated as an inhomogeneous Poisson process. At time step $\Delta t = 0.01\,\text{s}$, the rate $r_i$ of cell $i$ was

$$r_i(t) = g_i\,\omega_i(\mathbf{x}(t)) + b, \qquad b = 0.2\,\text{Hz}, \tag{4}$$

where $g_i$ is the gain (described below), $\omega_i$ is the spatial template (described above), and $b$ denotes the spatially uniform background firing rate.

For each cell, a target expected spike count $N_i^*$ was sampled with replacement from the empirical spike-count bank. The gain $g_i$ was then chosen so that the expected spike count matched $N_i^*$, subject to $g_i \geq 0$:

$$g_i = \max\left(0,\ \frac{N_i^* - bN_\Delta \Delta t}{\Delta t \sum_{t=1}^{N_\Delta} \omega_i(\mathbf{x}(t))}\right), \qquad K_i(t) \sim \text{Poisson}(r_i(t)\Delta t)\,. \tag{5}$$

Here $N_\Delta$ is the number of trajectory time bins and $K_i(t)$ is the number of spikes generated in time bin $t$. No positional jitter was included: all spikes generated in a given time bin were assigned exactly to the sampled trajectory position $\mathbf{x}_t$. Target spike counts were high enough so that clipping never occurred.

##### S3.2 Cellwise linear-$\lambda$ benchmark

The benchmark described in this subsection is used to test whether the SAC and CNN estimators recover known single-cell grid scales across the range relevant to the experimental recordings.

The cellwise simulation bank contained 100,000 equally spaced target spacings spanning $\lambda \in [20, 185]$ cm, with three stochastic realizations per target, yielding 300,000 synthetic cells. Realizations differed in orientation, phase, field width, spike-count target, field amplitudes, and Poisson noise (Supplementary Table 1), while sharing the trajectory and generative model described above. Cells were randomly divided into 240,000 training and 60,000 held-out examples. The held-out set therefore tests new stochastic realizations from the same trajectory and simulation distribution. Performance statistics were calculated using all 60,000 held-out cells; Supplementary Fig. 2a,c,d displays a random 30,000-cell subset for visual clarity.

**Supplementary Table 1.** Parameters varied in the cellwise synthetic benchmark.

| Quantity | Sampling rule |
|---|---|
| Grid spacing | 100,000 values uniformly spanning 20–185 cm; three stochastic realizations per value |
| Orientation | $\theta \sim \text{Uniform}(0°, 60°)$ |
| Relative phase | $r_x, r_y \sim \text{Uniform}(-1, 1)$ and $(x_c, y_c) = (\lambda r_x, \lambda r_y)$ |
| Field width | $\sigma \sim \text{TruncNormal}(0.13\lambda, 0.04\lambda; 0.05\lambda, 0.25\lambda)$ |
| Field amplitude | $A_{ik} \sim \text{TruncNormal}(1.0, 0.6; 0.15, 5.0)$, independently across fields |
| Spike count and trajectory | Target expected spike counts were sampled with replacement from retained module-1 (source publication indexing) cells in the Gardner rat-R day-1 reference recording; the trajectory was taken from the same source recording |
| Observation noise | Inhomogeneous Poisson sampling with $b = 0.2$ Hz and $\Delta t = 0.01$ s |

*Note.* $X \sim \text{TruncNormal}(\mu, s; L, U)$ denotes a normal distribution $\mathcal{N}(\mu, s^2)$ conditioned on $L \leq X \leq U$ and renormalized to integrate to unity. Thus, $\mu$ and $s$ are the location and standard deviation of the underlying untruncated normal distribution, whereas $L$ and $U$ are the lower and upper truncation bounds, respectively.

Except for field width, the simulated nuisance variables were sampled independently of target spacing and therefore could not serve as deterministic surrogate labels for $\lambda$. Field width increased with spacing on average but retained substantial variability at each target, making it informative without uniquely determining $\lambda$. The benchmark consequently evaluates whether the SAC or CNN can recover spacing from spatial structure under broad variation. It is not intended to capture all deformations or non-grid features observed in biological rate maps. Three exemplary rate maps built from the synthetic data are displayed in Supplementary Fig. 2b.

#### S3.3 Three-module benchmarks

The benchmarks described in this subsection are used to test whether the complete analysis pipeline preserves and distinguishes known constant- and decreasing-ratio module structures.

Two synthetic three-module populations were used to test recovery of the theory-discriminating ratio structure (Supplementary Fig. 3b,c). In both conditions, the largest- and smallest-scale modules were identical; only the middle-module distribution differed. This design isolates the local ratio structure while keeping the trajectory, observation model, module support, and outer scale distributions fixed.

The constant-ratio condition represents geometric module spacing of the kind predicted by the geometric-progression theories considered here. The selected ratio was close to $\sqrt{2}$ (Supplementary Table 2). The decreasing-ratio condition produced a larger ratio at the larger-scale transition than at the smaller-scale transition, as expected qualitatively under the peak-selection theory. Its requested arithmetic means gave $130/87 = 1.494$, $87/65 = 1.338$, and RoR $= 1.116$ (corresponding expected log-space values in Supplementary Table 2). These values closely approximate the $f = 0$ peak-selection predictions 3/2 and 4/3. The condition is a theory-relevant discrimination benchmark, not an exact realization of the full peak-selection model.

Within each module, cellwise spacings were lognormally distributed. This represents scale variation symmetrically in log space, where multiplicative differences become additive and reciprocal proportional deviations are treated equally. For a target arithmetic mean $m$ and standard deviation $s$,

$$\sigma_{\ln} = \sqrt{\ln\left(1 + \frac{s^2}{m^2}\right)}, \qquad \mu_{\ln} = \ln m - \frac{1}{2}\sigma_{\ln}^2, \qquad \lambda_i \sim \exp\left[\mathcal{N}(\mu_{\ln}, \sigma_{\ln}^2)\right]. \tag{6}$$

Because the downstream GMM is fitted in log scale, its back-transformed population component center is $\exp(\mu_{\ln})$, the geometric rather than arithmetic mean. Supplementary Table 2 therefore distinguishes ratios calculated from the requested arithmetic means from the expected ratios between the log-space component means.

**Supplementary Table 2.** Requested three-module scale distributions and corresponding benchmark targets. Module labels refer only to the largest (2), middle (3), and smallest (4) components of the synthetic triple and are not absolute anatomical module indices. Means and SDs are arithmetic values in centimeters. "Arithmetic RoR" is calculated from the requested arithmetic means; the final three columns give the expected ratios between the component centers fitted in log space, which are the quantities targeted by the analysis.

| Condition | $\lambda_2 \pm SD$ | $\lambda_3 \pm SD$ | $\lambda_4 \pm SD$ | Arithmetic RoR | $r_3^2$ | $r_4^3$ | Log-center RoR |
|---|---|---|---|---|---|---|---|
| Constant ratio | $130.0 \pm 8.0$ | $92.0 \pm 6.5$ | $65.0 \pm 2.2$ | 0.998 | 1.414 | 1.413 | 1.001 |
| decreasing ratio | $130.0 \pm 8.0$ | $87.0 \pm 6.5$ | $65.0 \pm 2.2$ | 1.116 | 1.496 | 1.336 | 1.120 |

The largest, middle, and smallest modules contained 200, 250, and 300 cells, respectively. All cells shared a fixed orientation of $\theta = 16°$, chosen as a representative oblique orientation that avoided exact alignment with the arena axes; the precise value was not assigned theoretical significance. Coherent module-specific orientations and systematic offsets from environmental boundaries have been observed experimentally.[3,4,18] Field width was fixed at $\sigma = 0.13\lambda$, chosen as a simple empirical approximation to the mean field-width–spacing relation in the Gardner rat-R day-1 maps used here.[12] For each synthetic cell, the phase and spike-count target were sampled independently with replacement from the empirical bank corresponding to its module. Spikes were then generated using the common Poisson observation model.

The generated populations were processed with the same rate-map construction, cellwise estimators, quality-control rules, logarithmic GMM fitting, component ordering, bootstrap-retention criteria, and ratio definitions as the experimental recordings.

Because these two three-module benchmarks use coherent, ideal hexagonal lattices under the same well-sampled trajectory, they test whether the analysis can preserve and distinguish known local ratio structures under controlled finite-arena observation noise. They do not establish robustness to limited module support, every deformation, module overlap, orientation dispersion, environmental remapping, or other feature of experimental grid-cell populations.

Cellwise estimator performance is reported in Stage 1 (Section S4). Recovery of the known three-module ratio structures by the complete downstream pipeline is reported in Stage 2 (Section S5).

### S4 Stage 1: Per-cell grid-scale estimation

#### S4.1 Spatial-autocorrelogram estimator

The spatial-autocorrelogram (SAC) estimator was used as the classical baseline for estimating grid scale and orientation. This estimator is standard in the grid-cell literature and was used in earlier reports of grid-module spacing values.[1,3] For each occupancy-corrected firing-rate map $r(x, y)$ of a grid cell, the SAC was computed at every integer spatial lag $\boldsymbol{\tau} = (\tau_x, \tau_y)$ as the Pearson correlation between the rate map and a translated copy of itself. The autocorrelogram was evaluated only over the mutually valid overlap

$$\Omega_{\boldsymbol{\tau}} = \{(x, y) : r(x, y) \text{ and } r(x + \tau_x, y + \tau_y) \text{ are finite}\} . \tag{7}$$

Because $\Omega_{\boldsymbol{\tau}}$ is recomputed for every displacement, the decreasing overlap at large spatial lags and near arena boundaries is handled explicitly rather than treated as zero-valued data.

The SAC value was then determined as follows:

$$C(\boldsymbol{\tau}) = \operatorname{corr}_{(x,y)\in\Omega_{\boldsymbol{\tau}}} \left(r(x, y), r(x + \tau_x, y + \tau_y)\right) . \tag{8}$$

Lags with fewer than 20 mutually valid bins were assigned NaN, thereby preventing correlations based on small overlaps near the SAC boundaries from contributing to peak detection. If the squared Pearson-denominator term was zero the corresponding SAC value was set to zero. For a rate map of dimensions $n_x \times n_y$, the resulting SAC had dimensions $(2n_x - 1) \times (2n_y - 1)$, with the zero-lag peak at its center.

For the biological datasets, the lag-wise Pearson correlations were evaluated using an implementation that explicitly accumulated the overlap count, sums, squared sums, and cross-products over the bins that were valid in both the original and shifted rate maps at each displacement. For the synthetic dataset with uniformly distributed grid scales (which contained 300,000 generated grid cells) and for the dataset with cropped experimental rate maps (described in the paragraph S4.2), an FFT-based implementation was used to reduce computational cost. In this implementation, linear cross-correlations of the finite-bin mask, rate map, and squared rate map were used to obtain the same lag-dependent moment terms. The direct and FFT backends, therefore, implemented the same NaN-aware Pearson SAC.

Autocorrelation fields were extracted from the SAC by first detecting local maxima with a nominal minimum spatial separation of 8 cm. This physical threshold was converted to the nearest integer number of SAC pixels for the spatial resolution used. With 1.5-cm bins, this yielded a minimum separation of 5 pixels under the Chebyshev metric, corresponding to 7.5 cm on the discretized SAC. The detected maxima were used as markers for watershed segmentation of the negated SAC, such that autocorrelation peaks acted as basins. Segmentation was restricted to pixels for which the original SAC value exceeded zero. Segmented fields were required to have a nominal minimum area of $0.015\,\mathrm{m}^2$. At the 1.5-cm bin resolution, this threshold corresponded to 67 pixels, or an effective discretized area of $0.015075\,\mathrm{m}^2$; smaller fields were discarded. The field containing the zero-lag position was subsequently removed. Field positions were represented by their local-maximum coordinates rather than by their centers of mass. The six remaining fields nearest to the SAC center were selected as the putative first ring of the hexagonal autocorrelation pattern.

Three grid axes were extracted from these six fields. The first axis was defined by the field direction closest to the horizontal direction; the other two were the nearest counter-clockwise and clockwise neighbours of this direction. When watershed segmentation split a single autocorrelation field and produced two nearly coincident axes, a geometric recovery procedure searched for a replacement field near the expected missing axis direction, approximately 60° from an intact axis, and at a comparable radial distance from the SAC center. Grid scale was then defined as the arithmetic mean of the three extracted axis lengths,

$$\lambda_{\mathrm{SAC}} = \frac{1}{3}\sum_{k=1}^{3} d_k, \tag{9}$$

where $d_k$ is the distance between the SAC center and the local maximum of the $k$-th selected autocorrelation field. Grid orientation was calculated as the circular mean of the three directed axis angles,

$$\theta_{\mathrm{SAC}} = \operatorname{atan2}\left(\sum_{k=1}^{3} \sin\theta_k, \sum_{k=1}^{3} \cos\theta_k\right). \tag{10}$$

SAC-derived estimates were subsequently subjected to cell-level quality control. A valid estimate required at least six off-center autocorrelation fields. In addition, the three adjacent inter-axis angles, after accounting for the 180° symmetry of each grid axis, were required to be strictly greater than 30° and strictly smaller than 90°. All pairwise ratios of the three axis lengths were required to lie between 0.5 and 2, and the cell was required to have a grid score of at least 0.2. These criteria determined whether an SAC-derived estimate was retained for subsequent module analysis.

In the implemented pipeline, SAC inputs are the same rate maps that are later passed to the CNN.

#### S4.2 Convolutional neural-network estimator

A convolutional neural network (CNN) was used as the second per-cell estimator because the synthetic benchmark showed that the SAC estimator rejected progressively more cells and increasingly underestimated spacing as grid scale approached the dimensions of the arena (Supplementary

Fig. 2a). For each grid cell, the CNN received only the rate-map-derived tensor described in the next paragraph. No recording, animal, or cell identifiers, module assignments, or population-level scale statistics were supplied as model inputs. Predictions were generated separately for individual cells. Grid-module membership and all between-module quantities were determined only after inference.

**Input representation and targets.** The source image was the same occupancy-normalized, edge-corrected rate map (Section S2) used for the SAC analysis. For CNN preprocessing, NaN bins were set to zero in the rate-map channel $r_{\text{norm}}$ and recorded in a binary visited-bin mask $M_{\text{visited}}$. The zero-filled rate map was divided by its maximum value, with a denominator floor of $10^{-6}$. The normalized rate maps were resized to $128 \times 128$ pixels by bilinear interpolation. The visited-bin masks were resized by nearest-neighbour interpolation and re-binarized.

The four input channels, in order, were

$$[r_{\text{norm}},\ M_{\text{visited}},\ X,\ Y]\,, \tag{11}$$

where $X$ and $Y$ were fixed CoordConv[19] coordinate channels spanning $[-1, 1]$ in the horizontal and vertical directions, respectively. These coordinate channels allowed the network to represent position relative to the map boundary. The physical window size, crop position, recording identity, and an indicator distinguishing full maps from crops were not input channels.

The CNN was trained jointly on grid scale and orientation so that its shared representation was constrained by both geometric properties of the lattice. For a square rate map with physical side length $L$, the fractional grid-scale target was

$$y_\lambda = \frac{\lambda}{1.5L}. \tag{12}$$

The value of $L$ was used only to construct the training target and to convert the output back to metres. It was not supplied to the CNN. Grid orientation was represented with its 60° periodicity as

$$\mathbf{y}_\theta = (\cos 6\theta,\ \sin 6\theta)\,. \tag{13}$$

**Network architecture and probabilistic outputs.** The model was a dual-head, multi-task CNN with a shared three-stage ConvNeXt-style encoder[20]. The base width was 32 channels, and the complete network contained 429,668 trainable parameters. A $4 \times 4$ convolution with stride 2 and padding 1 mapped the four input channels to 32 feature channels at a spatial resolution of $64 \times 64$, after which channel-wise layer normalization was applied independently at each spatial location. The three encoder stages had widths of 32, 64, and 128 channels and spatial resolutions of $64 \times 64$, $32 \times 32$, and $16 \times 16$, respectively. Each stage contained two residual ConvNeXt-style blocks. Each block comprised a $7 \times 7$ depthwise convolution, channel-wise layer normalization, a $1 \times 1$ pointwise expansion to four times the stage width, a Gaussian error linear unit (GELU) activation[21], and a $1 \times 1$ projection to the original width. Downsampling at the two transitions between stages was performed using $2 \times 2$ convolutions with stride 2. Neither dropout nor batch normalization was used.

Global average pooling produced a 128-dimensional feature vector. Separate scale and orientation heads each comprised a linear transformation from 128 to 64 features, a GELU activation, and a two-output linear layer.

If $(a_i, b_i)$ denotes the scale-head output for cell $i$, the predicted fractional mean and standard deviation were

$$\mu_{\lambda,i} = \mathrm{sigmoid}(a_i), \qquad \sigma_{\lambda,i} = \mathrm{softplus}(b_i) + 10^{-6}. \tag{14}$$

The physical grid-scale estimate and predicted standard deviation, both expressed in metres, were

$$\widehat{\lambda}_i = 1.5 L_i \mu_{\lambda,i}, \qquad \widehat{\sigma}_{\lambda,i} = 1.5 L_i \sigma_{\lambda,i}. \tag{15}$$

Here, $\widehat{\sigma}_{\lambda,i}$ is the conditional standard deviation of the Gaussian output distribution. It provides a model-based estimate of heteroscedastic aleatoric uncertainty, but does not separately quantify epistemic uncertainty.

The unconstrained orientation-head output for cell $i$, $\mathbf{v}_i = (v_{x,i}, v_{y,i})$, parameterized a von Mises distribution in $6\theta$ space. The orientation point estimate and the concentration parameter were

$$\widehat{\theta}_i = \frac{1}{6}\,\mathrm{atan2}(v_{y,i}, v_{x,i}) \pmod{\pi/3}, \qquad \kappa_i = \|\mathbf{v}_i\|_2. \tag{16}$$

**Phase 1: Synthetic pretraining.** Phase 1 used 300,000 simulated rate maps with known scale and orientation (Section S3). The maps were randomly divided into 240,000 training and 60,000 validation samples. The validation maps were not used for gradient updates. Training ran for 58 epochs with batch size 256 and AdamW.[22] The base learning rate was $1.6 \times 10^{-3}$ and weight decay was $10^{-4}$. The learning rate increased linearly from 1% of its base value during the first eight epochs (warm-up) and subsequently followed cosine decay. The global gradient norm was clipped at 5. The epoch-58 checkpoint minimized the sum of the scale and orientation validation negative log likelihoods and initialized Phase 2.

**Phase 2: Biological fine-tuning with synthetic anchors.** This second training phase was used to familiarize the CNN with biological variability and sampling noise that affect experimental rate maps in ways not fully represented by the synthetic data. Phase 2 fine-tuned the pretrained network using recordings from Vollan et al., together with a subset of the Phase-1 synthetic training maps retained as anchors. All targets for the biological samples originated from the SAC estimates computed from the corresponding full-arena rate maps. Biological samples were eligible for Phase-2 supervision only if they had a valid full-map SAC estimate, passed all SAC quality-control criteria described above, and had an SAC-estimated grid scale strictly below 1.05 m. This upper bound restricted the SAC pseudo-labels to the range below the large-scale regime in which the synthetic benchmark showed that the SAC estimator increasingly underestimated grid spacing.

**Biological crop construction.** Full-map samples represented the complete 1.5 m × 1.5 m arena. To expose the CNN to biological rate-map texture containing fewer visible grid fields while retaining reliable scale and orientation labels estimated from the full arena, we additionally generated 1.05 m × 1.05 m crops. A cropped rate map with a relatively small physical grid scale contains fewer complete lattice fields than the corresponding full-arena rate map and therefore approximates, in fractional terms, the limited field visibility encountered for larger grid scales.

All full maps and crops were resized to 128 × 128 pixels. Neither the physical window size $L$ nor an indicator distinguishing full maps from crops was supplied to the CNN. The network therefore could not determine the physical size of the sampled region from the tensor dimensions, and the prediction target represented grid scale relative to the sampled window. Because the scale target was normalized by map width (Eq. 12), a given physical scale occupied a larger fraction of a crop than of the full arena. The crops therefore extended biological training coverage towards the fractional-scale regime characteristic of cells with large grid scales while retaining labels estimated from more informative full-arena rate maps. For example, a 150-cm grid scale represented in a 150-cm full map and a 105-cm grid scale represented in a 105-cm crop both correspond to a target $y_\lambda = 2/3$.

Crop rate maps were not produced by cutting regions from the already smoothed full-arena rate maps. Instead, they were reconstructed from the underlying spike-count and occupancy histograms. A 1.05 m $\times$ 1.05 m crop window was moved across the full arena in steps of three 1.5-cm spatial bins. This produced an 11 $\times$ 11 grid of window positions, or 121 crops per eligible cell. For each window, the spike-count and occupancy histograms were cropped first and then smoothed separately with a Gaussian kernel of 4-cm standard deviation. Both smoothed histograms were corrected for kernel support truncated at the crop boundary. Bins that did not exceed the occupancy threshold derived from the corresponding full-arena data were masked, and the crop rate map was obtained by dividing the corrected, smoothed spike-count map by the corrected, smoothed occupancy map.

Each crop inherited the grid scale and orientation labels estimated by the SAC method from the corresponding cell's full-arena rate map. Crop position was not supplied to the CNN. Although cropping increased the representation of biological rate maps containing few complete grid fields, an analytical crop edge is not equivalent to a physical wall, near which the animal's movement and spatial sampling may differ.

**Animal-level data partition.** The biological recordings were partitioned by the animal identifier preceding the underscore in the recording name. All recordings from an animal were assigned to one split:

- Training (9 animals and 17 recordings): `25691_1`, `25691_2`; `25953_4`, `25953_5`; `26034_3`; `26035_1`; `26648_1`, `26648_2`; `27764_1`; `27765_1`, `27765_2`, `27765_3`; `28063_1`, `28063_4`, `28063_5`; and `28229_2`, `28229_3`.
- Validation (4 animals and 6 recordings): `24365_2`; `28258_4`; `28304_1`, `28304_2`; and `29502_1`, `29502_3`.
- Test (2 animals and 3 recordings): `25127_1`; and `25843_1`, `25843_2`.

No animal in the biological training split contributed to the RoR analysis reported in the Main Text. Recording `26018_2` was excluded from these splits because the arena used there was apparently of a circular shape.

The biological training pool contained 2,429 full maps and 293,909 crops, for a total of 296,338 biological samples. The validation and test sets contained 693 and 730 full maps, respectively. The union of the validation and test recordings additionally yielded 172,183 evaluation crops. These crops were used only for diagnostic evaluation and not for gradient updates.

**Synthetic anchors and Phase-2 sampling.** Phase 2 additionally contained 98,681 synthetic anchors uniformly drawn without replacement from the Phase-1 training pool. The complete stored Phase-2 pool therefore contained 395,019 samples.

A weighted random sampler drew samples with replacement and requested 395,019 indices per epoch. The sampler was constructed to assign source-level probability masses of 0.15, 0.45, and 0.40 to biological full-arena rate maps, biological crops, and synthetic anchors, respectively. Within each of the two biological sources, the fractional scale distribution was balanced using adjacent, non-overlapping, half-open intervals of width 0.01. Equal conditional probability mass was assigned to every populated interval within each biological source.

Positive per-sample weights were normalized to have mean one and were capped at 50 to limit the influence of sparsely populated intervals. Because of this cap, the realized source proportions could differ slightly from the nominal 0.15/0.45/0.40 targets. They also varied stochastically between epochs because sampling was performed with replacement.

**Training objectives and GradNorm.** For a batch of $N$ samples, let $y_{\lambda,i}$ denote the dimensionless scale target, $\mathbf{y}_{\theta,i}$ the unit-vector orientation target, and let $\mu_{\lambda,i}$, $\sigma_{\lambda,i}$, and $\mathbf{v}_i$ denote the corresponding network outputs.

During the first eight epochs of each training phase, deterministic objectives were used to establish stable point predictions. The scale loss was given by the mean-squared error,

$$\mathcal{L}_\lambda^{\text{warm}} = \frac{1}{N}\sum_{i=1}^{N} (\mu_{\lambda,i} - y_{\lambda,i})^2 \,, \tag{17}$$

and the orientation loss was the cosine distance,

$$\mathcal{L}_\theta^{\text{warm}} = \frac{1}{N}\sum_{i=1}^{N} \left[ 1 - \frac{\mathbf{v}_i^{\mathsf{T}} \mathbf{y}_{\theta,i}}{\|\mathbf{v}_i\|_2 \|\mathbf{y}_{\theta,i}\|_2} \right] . \tag{18}$$

Because $\mathbf{y}_{\theta,i}$ is a unit vector, its norm is one, but is retained in the expression for completeness.

From epoch 9 onward, the scale output was trained by Gaussian negative log likelihood,

$$\mathcal{L}_\lambda^{\text{NLL}} = \frac{1}{N}\sum_{i=1}^{N} \frac{1}{2} \left[ \left( \frac{y_{\lambda,i} - \mu_{\lambda,i}}{\widetilde{\sigma}_{\lambda,i}} \right)^2 + 2 \ln \widetilde{\sigma}_{\lambda,i} + \ln(2\pi) \right] , \tag{19}$$

where $\widetilde{\sigma}_{\lambda,i} = \max(\sigma_{\lambda,i}, 10^{-4})$ was used for numerical stability during training.

Orientation was optimized with the von Mises negative log likelihood in $6\theta$ space:

$$\mathcal{L}_\theta^{\text{NLL}} = \frac{1}{N}\sum_{i=1}^{N} \left[ -\mathbf{v}_i^{\mathsf{T}} \mathbf{y}_{\theta,i} + \ln I_0^e(\kappa_i) + \kappa_i \right] , \tag{20}$$

where $I_0^{\text{e}}(\kappa) = \exp(-\kappa) I_0(\kappa)$ is the exponentially scaled modified Bessel function of the first kind. In equation 20, the additive constant $\ln(2\pi)$ is omitted, which does not affect optimization.

To dynamically balance the scale and orientation objectives, their task weights were adapted using a first-order implementation of GradNorm.[23] The positive weights $w_\lambda$ and $w_\theta$ were normalized after each update such that $w_\lambda + w_\theta = 2$. GradNorm used the shared encoder parameters to calculate task gradient norms, an asymmetry exponent $\alpha = 0.5$, and a separate Adam optimizer with learning rate $10^{-3}$. The GradNorm weights were updated every second minibatch and were reset when training changed from the deterministic objectives to the likelihood objectives at epoch 9.

From epoch 9 onwards, each likelihood objective supplied to GradNorm was shifted according to

$$\widetilde{\mathcal{L}}_j = \mathcal{L}_j - \text{stopgrad}(b_j) + 10^{-3}, \qquad j \in \{\lambda, \theta\}, \tag{21}$$

where $b_j$ was a task-specific normalization term evaluated over the current minibatch and stopgrad denotes detachment from automatic differentiation. For the scale objective, $b_\lambda$ contained the Gaussian normalization contribution $\ln \sigma_\lambda + \frac{1}{2}\ln(2\pi)$; for the orientation objective, $b_\theta$ contained the von Mises contribution $\ln I_0^e(\kappa)$. Subtracting these terms produced non-negative excess-loss values, thereby stabilizing the relative inverse training rates used by GradNorm to update the task weights. Because the subtracted terms were detached,

$$\nabla_{\mathbf{w}} \widetilde{\mathcal{L}}_j = \nabla_{\mathbf{w}} \mathcal{L}_j \tag{22}$$

for the CNN parameters $\mathbf{w}$. The shift therefore changed the numerical task-loss values used by GradNorm without changing the corresponding CNN-parameter gradients.

During the initial synthetic pretraining (Phase 1), GradNorm balanced the shifted scale and orientation objectives over the synthetic samples. During the biological fine-tuning (Phase 2), GradNorm was restricted to the biological objectives, whereas the synthetic losses bypassed GradNorm and entered as a unit-weight anchor to preserve performance on that part of the synthetic domain where the grid scales did not feature in the experimental samples. From epoch 9 onwards, the final Phase-2 objective was therefore

$$\mathcal{L}_{\text{phase 2}} = w_\lambda \widetilde{\mathcal{L}}_{\lambda,\text{bio}} + w_\theta \widetilde{\mathcal{L}}_{\theta,\text{bio}} + \eta \left(\mathcal{L}_{\lambda,\text{syn}} + \mathcal{L}_{\theta,\text{syn}}\right), \qquad \eta = 1. \tag{23}$$

**Augmentation, optimization, and model selection.** During both phases, every training item was subjected to a random rotation by $k \times 90°$ degrees, where $k$ was sampled uniformly from $\{0, 1, 2, 3\}$, followed independently by a horizontal reflection with probability 0.5. The visited mask and orientation target were transformed consistently with the rate map. Validation and test maps were not augmented.

Phase 2 ran for 58 epochs with batch size 256, AdamW, base learning rate $2 \times 10^{-4}$, weight decay $10^{-4}$, and the same eight-epoch linear-warm-up and subsequent cosine-decay schedule as Phase 1. The global gradient norm was clipped at 5. During the first eight Phase-2 epochs, the convolutional stem and first encoder stage were frozen; all parameters were trainable after that.

The Phase-2 checkpoint used in the reported analyses was epoch 25, selected by minimizing the sum of the scale and orientation negative log likelihoods on the biological full-map validation split. The biological test split and evaluation crops were not used for checkpoint selection. On the held-out biological full-map test split ($n = 730$ cells), the CNN agreed closely with the SAC-derived reference labels, with a grid-scale mean absolute error (MAE) of 2.76 cm, a root-mean-square error (RMSE) of 5.32 cm, and an orientation MAE of 1.41°. The median absolute orientation error was 0.75°.

On the 60,000 synthetic validation maps, for which scale and orientation were known from the data-generating process, grid-scale MAE and RMSE were 1.37 cm and 3.27 cm, respectively, and orientation MAE and median absolute orientation error were 0.86° and 0.12°, respectively (Supplementary Fig. 2c–d). The synthetic validation split was not used for gradient updates, but was used to select the Phase-1 checkpoint.

On the diagnostic biological crop evaluation set ($n = 172{,}183$ crops), grid-scale MAE and RMSE were 4.08 cm and 6.34 cm, respectively, and orientation MAE and median absolute orientation error were 2.49° and 1.51°, respectively (Supplementary Fig. 2f).

Because the biological reference values were obtained from SAC estimates, the biological evaluation quantifies agreement with SAC within the reliable pseudo-label range, whereas the synthetic evaluation provides the direct assessment against known ground truth.

**Inference and downstream use.** For the reported module analysis, the selected model was applied independently to every full rate map. The CNN branch did not require the SAC peak-count, inter-axis-angle, or axis-length-ratio criteria. The SAC nevertheless supplied the gridness criterion: cells with a stored grid score below 0.2 were excluded. The final module-analysis did not impose a hard threshold on the uncertainty predicted by the selected CNN. Instead, the predicted per-cell scale standard deviations were retained for uncertainty-aware population inference.

Grid-module inference was performed on $x_i = \log_{10} \widehat{\lambda}_i$. The CNN scale standard deviation was transformed to log-scale units with the first-order delta method,

$$\tau_i = \frac{\widehat{\sigma}_{\lambda,i}}{\widehat{\lambda}_i \ln 10}. \tag{24}$$

The pairs $(x_i, \tau_i)$ were supplied to both the heteroscedastic KDE and the heteroscedastic GMM, including their full-data fits and bootstrap refits (see section S5.3). During bootstrap resampling, point estimates and their corresponding predicted uncertainties were resampled jointly. The SAC branch had no corresponding cell-specific standard deviation estimate and therefore used the standard KDE and GMM procedures.

#### S4.3 Geometric field-center estimator

As an explicit alternative to autocorrelogram-based scale estimation (SAC method), we also built and evaluated a geometric field-center pipeline based on boundary-aware multiscale Laplacian-of-Gaussian detection. The rationale was that, if the main limitation of the SAC estimator at large grid scales arose from inward-biased peak locations in the autocorrelogram, driven by truncated fields at the arena boundary, then a more direct analysis in the original rate map – localizing putative field centers first and estimating spacing only afterwards – might recover scale more faithfully while remaining interpretable. Candidate fields were therefore detected across scales, refined locally, and converted into a spacing estimate using either robust nearest-neighbor distances or a global lattice-based fit, with explicit handling of boundary-truncated fields, including fits whose inferred centers lay outside the sampled arena. Individual per-cell estimates can also be weighted in this method.

In practice, however, this did not remove the main large-scale failure mode. One problem is that, if spacing is estimated only from well-contained interior fields, performance must deteriorate at large $\lambda$, because the number of fully visible fields falls rapidly with scale,

$$N_{\text{vis}} \propto \frac{A_{\text{arena}}}{\lambda^2},$$

and the remaining retained centers are no longer representative of the full lattice. More importantly, allowing truncated fields into the estimator does not resolve this limitation cleanly. When only a partial field is observed, center location and field width must be inferred jointly from incomplete and noisy support so the local fit becomes weakly identifiable. Under conservative settings, the detector therefore often abstained because too few reliable centers remained for a stable spacing estimate. Under more permissive settings, yield increased, but the added centers were disproportionately drawn from partial or weakly resolved edge fields, which compressed nearest-neighbor distances and biased lattice fits toward smaller spacings. Thus, the large-scale regime imposed a fundamental trade-off between retention and accuracy rather than yielding a clear geometric rescue of SAC.

For this reason, we treat the blob-based estimator as an important methodological control rather than a viable primary solution. It shows that the large-scale limitation is not specific to autocorrelograms, but reflects the finite-field, finite-arena geometry of the problem itself. A center-detection approach may remain useful for visualization or sanity checks, but is unlikely to systematically outperform SAC in the largest-scale regime, because both methods ultimately rely on geometric information that becomes intrinsically underconstrained once only fragments of the lattice are observed.

The key failure mode for the boundary-refit branch was not merely boundary truncation per se, but an identifiability problem in the truncated local fit. In the simplest 1D analog, if only one flank of an approximately Gaussian field is sampled, then fitting both center $\mu$ and width $\sigma$ produces an extended ridge in parameter space: shifts in $\mu$ can be partly compensated by changes in $\sigma$ while preserving a very similar likelihood over the observed support. In practice, finite trajectory sampling, rate-map smoothing, and spike-count noise broaden this ridge further. The result is a high-variance and often biased estimate of field center for strongly truncated fields, even when the fitted model class is otherwise appropriate. This explains why explicitly permitting centers outside the arena was necessary but not sufficient: once truncation is too severe, the inferred center is no longer geometrically well constrained, and later spacing estimates inherit that instability.

### S5 Stage 2: module-level scale estimation

Stage 2 converted the retained cellwise scale estimates into recording-level module centers. The procedure was applied separately to each recording and Stage-1 estimator. SAC-derived scales entered this analysis only after fulfilling the SAC geometric quality-control criteria described above. CNN-derived scales were retained when the cell's SAC grid score was at least 0.2, while the other SAC quality-control criteria were not imposed on this branch. No hard threshold was applied to the CNN-predicted scale uncertainty. Lattice orientation was not used to define the grid modules.

All scale clustering was performed in logarithmic coordinates,

$$x_i = \log_{10}\left(\frac{\widehat{\lambda}_i}{1\,\mathrm{m}}\right), \tag{25}$$

because multiplicative scale ratios become differences between module centers. In this section, $x_i$ is the log-transformed scale estimate of cell $i$. If $\mu_m$ and $\mu_{m+1}$ are two GMM-component centers in log space, their adjacent ratio is

$$r^{m}_{m+1} = 10^{\mu_m - \mu_{m+1}}. \tag{26}$$

Thus, a constant-ratio sequence has equally spaced centers in log space.

#### S5.1 Kernel-density estimate

For a recording containing $n$ retained cells, the standard spacing kernel density estimate (KDE) was

$$\widehat{f}(x) = \frac{1}{n}\sum_{i=1}^{n} \frac{1}{\sqrt{2\pi h^2}} \exp\left[-\frac{(x-x_i)^2}{2h^2}\right], \qquad h = 0.065\,n^{-1/5}, \tag{27}$$

where $h$ is expressed in $\log_{10}(\lambda/\mathrm{m})$ units. The density was evaluated at 1,000 equally spaced points over the interval from $\min(0.30\,\mathrm{m}, 0.9\lambda_{\min})$ to $\max(1.80\,\mathrm{m}, 1.1\lambda_{\max})$, thereby covering at least 0.30–

1.80 m while adding a 10% margin when the padded observed range extended beyond these limits. Candidate modules were local density maxima detected with `scipy.signal.find_peaks`. In the KDE, the minimum-height setting was 0.02 of the maximum density attainable if all kernels were centered at the same position, and the minimum prominence was 0.0005.

The number $K$ of detected KDE maxima set the number of Gaussian-mixture-model (GMM) components. Consequently, the KDE and GMM summaries are related rather than independent: KDE peaks provide both the component count and the initialization for component means.

#### S5.2 Gaussian-mixture estimate

For the SAC branch, the primary module centers were obtained from a Gaussian mixture model (GMM) fitted to the same $x_i$ values (Eq. 25),

$$p(x_i) = \sum_{k=1}^{K} \pi_k \mathcal{N}(x_i \mid \mu_k, v_k), \qquad \pi_k \geq 0, \qquad \sum_{k=1}^{K} \pi_k = 1. \tag{28}$$

Here, $i = 1, \ldots, n$ indexes cells, and $k = 1, \ldots, K$ indexes the GMM components. The number of components $K$ was set to the number of KDE maxima. The term $\mathcal{N}(x_i \mid \mu_k, v_k)$ denotes the Gaussian probability density of component $k$, with mean $\mu_k$ and variance $v_k$ in log-scale units, and $\pi_k$ is its mixture weight.

The standard model was fitted using the expectation-maximisation algorithm implemented in `GaussianMixture` from `sklearn.mixture`. GMM-component means were initialized at the detected KDE maxima. Mixture weights, means, and component variances were estimated from the data. Each cell was assigned to the component with the largest fitted posterior probability. Components were then ordered by their means and their scale representatives were obtained as $10^{\mu_k}$ meters. All peak-count-eligible standard-GMM bootstrap refits converged, the maximum iteration count was 55, below the limit of 100 iterations. Only recordings with three KDE-peaks and, therefore, three GMM-components entered the subsequent adjacent-ratio and ratio-of-ratios analyses.

#### S5.3 Propagation of CNN predictive uncertainty

The SAC estimator did not provide an analogous per-cell predictive standard deviation. The SAC branch therefore used the standard versions of the KDE (Eq. 27) and GMM (Eq. 28) in both the full-data fit (i.e., all cells retained after the SAC filtering) and the cell bootstrap. For the CNN branch, the predicted scale standard deviation in meters $\widehat{\sigma}_{\lambda,i}$ (Eq. 15) was transformed to log units by the first-order delta method,

$$\tau_i = \frac{\widehat{\sigma}_{\lambda,i}}{\widehat{\lambda}_i \ln 10}. \tag{29}$$

The CNN KDE replaced the common kernel variance in Eq. (27) by $h^2 + \tau_i^2$,

$$\widehat{f}_{\mathrm{CNN}}(x) = \frac{1}{n} \sum_{i=1}^{n} \frac{1}{\sqrt{2\pi(h^2 + \tau_i^2)}} \exp\left[-\frac{(x - x_i)^2}{2(h^2 + \tau_i^2)}\right]. \tag{30}$$

Thus, an uncertain prediction contributed a broader, lower kernel rather than being removed or assigned an explicit inverse-variance weight.

Then the full-data GMM, used to obtain the raw grid-module centers and cell assignments, was initialized by the uncertainty-aware KDE maxima. CNN uncertainty was also incorporated into both the full-data GMM and its bootstrap refits through the heteroskedastic measurement-error mixture

$$p(x_i \mid \tau_i) = \sum_{k=1}^{K} \pi_k \, \mathcal{N}\Big(x_i \,\Big|\, \mu_k, v_k + \tau_i^2\Big) \,. \tag{31}$$

This model is a specialization of an error-deconvolving Gaussian mixture model for observations with heterogeneous, cell-specific uncertainty.[24] Here, $v_k$ denotes the intrinsic within-component variance, whereas $\tau_i^2$ is the estimated cell-specific observation variance, treated as fixed during GMM fitting. Consequently, for cells with larger CNN-predicted scale uncertainty, the Gaussian density associated with each GMM component was broader. The model was fitted with a custom expectation-maximisation algorithm. For every full-data or bootstrap fit, component means were initialized at the corresponding KDE maxima, weights were initialized uniformly, and intrinsic component variances were initialized to the variance of the fitted $x_i$ values divided by $K^2$.

A heteroskedastic fit was considered converged when the absolute change in the total log-likelihood was below $10^{-4}$, with a maximum of 1,000 iterations. Non-converged heteroskedastic bootstrap refits were excluded. All full-data heteroskedastic GMMs converged.

#### S5.4 Recording-level cell bootstrap

For each estimator and recording, 5,000 bootstrap samples of size $n$ were drawn with replacement from the retained cells. CNN scale estimates and their predicted standard deviations were resampled as pairs; scale coordinates were not additionally jittered. The spacing KDE was recomputed on the full-data evaluation grid, candidate modes were redetected, and the GMM was refitted in every replicate. SAC replicates used the standard versions of KDE and GMM, whereas CNN replicates used the adaptive heteroskedastic versions of KDE (Eq. 30) and GMM (Eq. 31). A replicate contributed to the module-center and ratio distributions only when it recovered the same number of KDE modes as the full-data fit; the CNN GMM was additionally required to converge. Replicate KDE peaks and sorted GMM means were matched one-to-one to the nearest reference positions in full data. The adjacent-module ratios and the ratio-of-ratios (RoR) were then recomputed in the full-data module order. Bootstrap confidence 95% percentile intervals were reported.

The primary empirical pipelines are SAC+GMM and CNN+GMM. The SAC+KDE and CNN+KDE pipelines retain the same upstream cell estimates and uncertainty treatment but use the KDE maxima as the module representatives, and serve as robustness analyses. The empirical module centers are shown in Supplementary Fig. 1c–d. The module-assignment logic and synthetic validation are shown in Supplementary Fig. 3, where in panels b,c one can see a comparison of all four pipelines on the two synthetic benchmarks that featured three grid modules. For the SAC estimator, KDE yielded a significantly lower bootstrap distribution of RoR values compared to the GMM's distribution: ($p = 0.016$) on the constant-ratio dataset and ($p = 0.009$) on the decreasing-ratio dataset. For the CNN estimator, the difference between the KDE and GMM distributions was not significant on the constant-ratio dataset ($p = 0.882$) and on the decreasing-ratio dataset ($p = 0.252$). Based on these benchmark results, GMM was chosen as a primary method for the module-level scale estimation.

### S6 Ratios, recording bootstrap, within-animal Monte Carlo combination, and population statistics

This section defines the ratios of interest, carries conditional cell-resampling uncertainty into the within-animal combinations and reliability weights, and defines the animal-level population summaries and comparisons with theoretical references.

#### S6.1 Ratio definitions and logarithmic working scale

This subsection defines the two adjacent scale ratios and their ratio of ratios, and explains why statistical calculations were performed on a logarithmic scale.

Using the module order defined in the main text, and the conditional peak-selection indexing summarized in Section S1,

$$r_3^2 = \frac{\lambda_2}{\lambda_3}, \qquad r_4^3 = \frac{\lambda_3}{\lambda_4}, \qquad \mathrm{RoR}_4^2 = \frac{r_3^2}{r_4^3} = \frac{\lambda_2\lambda_4}{\lambda_3^2}. \tag{32}$$

Thus, unity (RoR = 1) means that the three module centers are equally spaced on a logarithmic scale, or equivalently that $\lambda_3$ is the geometric mean of $\lambda_2$ and $\lambda_4$. The conservative peak-selection reference for this triple is $49/45 \approx 1.089$, obtained for $f = 1/2$.[11]

The logarithmic treatment of the cell-scale distributions and the GMM is described in Section S5. After the fitted component centers were back-transformed to linear scale and the ratios were formed, each ratio or RoR value ($R$) was represented as $z = \ln R$ for the recording-, animal-, and population-level calculations. These quantities are multiplicative, so population centers, residuals, bootstrap flanks, reliability weights, intervals, and reference comparisons were calculated in log space and exponentiated for display.

Recording-level adjacent ratios and their primary population summaries are shown in Supplementary Fig. 4. RoR distributions, reference comparisons, and population summaries are shown in Supplementary Fig. 5. Supplementary Figure 1e–h compares these population summaries across alternative analysis units, weighting rules, and centering specifications.

#### S6.2 Recording-level cell bootstrap

For each recording, the retained cell-scale estimates were resampled with replacement 5,000 times. Every replicate was passed through the same log-space GMM fitting, component-ordering, and ratio-construction procedure as the full data. Module assignments and component centers were therefore refitted rather than held fixed, because uncertainty in the fitted module structure contributes to uncertainty in the ratios. A bootstrap replicate contributed to a three-module statistic only when the refitted analysis recovered the required three ordered, finite component centers and produced a finite statistic. The resulting bootstrap distributions are therefore conditional on successful three-module recovery. Retained counts range from 3,014 to 4,993 for SAC and from 3,732 to 5,000 for CNN and are reported for every recording in Supplementary Table 5.

The retained bootstrap propagates finite-cell sampling from the observed pool, the influence of individual cells, module-center fitting, and nonlinear ratio construction under the selected log-space GMM representation. It cannot recover cells excluded upstream, identify an unsampled tail or biological module, establish that the GMM is the correct biological distribution, capture variability arising from CNN retraining, random initialization, or computational nondeterminism, or diagnose a bias shared by the estimator across recordings. Accordingly, a narrow retained bootstrap interval demonstrates stability under resampling of the observed cell pool; it does not establish that a sparsely represented module was sampled without tail truncation or that its absolute center is free of scale-dependent estimator bias.

For a recording $i$ with raw (i.e., full-data, non-bootstrap) value $R_i$: $z_i = \ln R_i$, and $z_i^{(b)} = \ln R_i^{(b)}$ are the retained bootstrap values. The recording bootstrap was summarized by its median and central 95% quantiles,

$$m_i = \operatorname{median} z_i^{(b)}, \qquad \ell_i = m_i - q_{i,.025}, \qquad u_i = q_{i,.975} - m_i, \qquad h_i = \max(\ell_i, u_i). \tag{33}$$

The longer flank $h_i$ was used for reliability so that a skewed distribution was not assigned high precision based on its shorter tail. Here and below, reliability refers only to conditional precision under resampling of the observed cells and the fitted analysis pipeline; it is not a general measure of recording quality or biological importance.

The percentile position of the raw estimate within its retained bootstrap was

$$\pi_i = \frac{1}{B_i} \sum_{b=1}^{B_i} \mathbf{1}\left[z_i^{(b)} \leq z_i\right]. \tag{34}$$

Supplementary Table 5 reports $R_i$, the bootstrap median and central interval, retained count, and $\pi_i$ for every recording. The reliability weight depends only on $h_i$; the full table and recording-level distributions retain the additional information about asymmetry and the position of the raw estimate within the retained bootstrap distribution. The retained-replicate fraction was reported separately and did not enter the reliability weight.

The raw full-data estimate was used as the primary recording point because it was calculated from all retained cells in the recording. The bootstrap median was treated as a centering sensitivity rather than an automatic bias correction, because displacement can reflect nonlinearity, influential cells, skewness, or conditional replicate retention.

##### S6.3 Combination of repeated recordings within animal

This subsection describes the combination of repeated recordings within a given animal while propagating the conditional uncertainty represented by their recording-level bootstrap distributions.

Animals were the independent biological units, and recordings were nested within animals. Repeated recordings from the same animal represented distinct recording sessions; for Gardner rat `R`, the two analyzed recordings corresponded to day 1 and day 2.

Because repeated recordings differed in finite-cell support and bootstrap stability, the primary within-animal combination gave more influence to recordings with narrower full-pipeline RoR distributions. The weights used bootstrap width alone and contained neither the observed RoR nor either theoretical reference. Because the width and point estimate were nevertheless estimated from the same recording, the resulting weights need not be statistically independent of the observed values. Equal-weight analyses therefore assessed sensitivity to the reliability-weighting rule (see Supplementary Fig. 1e–h).

For each estimator and statistic separately, a common recording-level regularization floor was calculated across all included recordings:

$$\epsilon_{\text{rec}} = \text{median}_{i \in \mathcal{I}}(h_i), \tag{35}$$

where $\mathcal{I}$ is the full set of included recordings for that estimator and statistic. This floor was not recalculated separately within each animal. Recording weights were

$$w_i = \frac{1}{h_i^2 + \epsilon_{\text{rec}}^2}, \qquad \alpha_i = \frac{w_i}{\sum_{j \in S_a} w_j}, \tag{36}$$

where $S_a$ contains the recordings of animal $a$. The animal's raw point estimate was

$$y_a = \sum_{i \in S_a} \alpha_i z_i. \tag{37}$$

The median longer flank regularized the weights at the scale of a typical recording and prevented an exceptionally narrow bootstrap distribution from producing arbitrarily large leverage.

Recording-level uncertainty was propagated by 30,000 Monte Carlo draws. On every draw, one retained bootstrap value was sampled with replacement and independently from the retained bootstrap distribution of each recording belonging to the animal and combined using the fixed full-data recording weights:

$$Y_a^{(t)} = \sum_{i \in S_a} \alpha_i z_i^{(b_i(t))}, \qquad t = 1, \ldots, 30{,}000. \tag{38}$$

Thus every recording from animal $a$ contributes to every MC draw in proportion to its recording-level weight.

Independent sampling across recordings is a conditional approximation. It propagates the recorded bootstrap distributions (shape, skewness, and asymmetry) but does not represent additional independent biological replication and does not add a separate variance component for disagreement between sessions from one animal. Under this construction, combining repeated recordings may narrow the resulting animal-level distribution.

The median of the resulting animal-level MC distribution is $m_a$, and $L_a$ and $U_a$ denote its lower and upper central-95% flanks, with $H_a = \max(L_a, U_a)$. The width $H_a$ describes the conditional uncertainty of the combined animal estimate rather than the uncertainty of any one recording. For an animal represented by one recording, the same procedure reduces to sampling from that recording's retained bootstrap distribution.

#### S6.4 Population center and displayed summary objects

This subsection defines the animal-level reliability weights, the population center, and the three distinct uncertainty summaries displayed in the figures.

In the primary analysis, the population units were animals. In the recording-as-unit sensitivity described below, the same population formulas were applied directly to recordings. For animals represented by repeated recordings, $H_a$ was obtained only after their recording bootstrap distributions had been combined. The weighting therefore reflected the conditional precision of the animal-level estimate rather than that of any single recording.

For each estimator and statistic separately, the animal-level regularization floor and weights were

$$\epsilon_{\text{animal}} = \text{median}_a(H_a), \qquad v_a = \frac{1}{H_a^2 + \epsilon_{\text{animal}}^2}, \qquad \beta_a = \frac{v_a}{\sum_c v_c}. \tag{39}$$

The primary population center was the weighted mean of the raw animal estimates in natural-log space:

$$\mu = \sum_a \beta_a y_a, \qquad \widehat{R}_{\text{pop}} = \exp(\mu). \tag{40}$$

The regularization floors limited leverage and were not estimates of biological heterogeneity. Weight concentration was summarized by the effective sample size, $1/\sum_a \beta_a^2$, and by the maximum animal weight. Both are reported with the population results in Supplementary Table 6.

The figures display three distinct summary objects around the same reliability-weighted mean. These answer different questions: how much the observed animal estimates vary ("Spread"), how precisely the weighted population mean is estimated under the selected HC1 (heteroskedasticity-consistent) approximation ("95% CI"), and how much recording-level bootstrap uncertainty is typically propagated into an animal estimate ("MC flanks").

**Observed spread.** The bias-corrected weighted SD of the animal log estimates was

$$s_w = \left[\frac{\sum_a \beta_a (y_a - \mu)^2}{1 - \sum_a \beta_a^2}\right]^{1/2}. \tag{41}$$

The displayed range $\exp(\mu \pm s_w)$ describes the observed weighted variation among animal estimates, including remaining measurement error.

**HC1 confidence interval.** For $k$ independent animals, the HC1 standard error $\widehat{\text{SE}}_{\text{HC1}}$ of the weighted log-scale population center was

$$\widehat{\text{SE}}_{\text{HC1}} = \left[\frac{k}{k-1} \sum_a \beta_a^2 (y_a - \mu)^2\right]^{1/2}, \tag{42}$$

and the 95% interval was

$$\text{CI}_{95\%}^{\text{HC1}} = \left[\exp\left(\mu - t_{0.975,k-1}\widehat{\text{SE}}_{\text{HC1}}\right), \exp\left(\mu + t_{0.975,k-1}\widehat{\text{SE}}_{\text{HC1}}\right)\right]. \tag{43}$$

This is the approximate inferential interval used for comparisons with fixed references. The sandwich form allows the residual contribution to differ among animal estimates rather than imposing a common residual variance, and the $t_{k-1}$ multiplier provides a small-sample reference approximation[25–27]. For this interval and the corresponding reference comparisons, the selected reliability weights were treated as fixed. Because only five or eight animals were available and the reliability weights were estimated from the same data, coverage and reference comparisons are not exact finite-sample inference.

**Typical animal-level MC flanks.** Typical conditional uncertainty was summarized by

$$\widetilde{L} = \mathrm{median}_a(L_a), \qquad \widetilde{U} = \mathrm{median}_a(U_a), \tag{44}$$

and displayed as $[\exp(\mu - \widetilde{L}), \exp(\mu + \widetilde{U})]$. These medians were unweighted so that the display represents the uncertainty of a typical animal estimate rather than being dominated by the most precise animals. This is neither a population-dispersion interval nor a confidence interval for the population center.

#### S6.5 Reference comparisons

This subsection defines the approximate HC1–$t$ comparisons of the population center with unity and with the conservative peak-selection reference 49/45.

For the unity and peak-selection references $r \in \{1, 49/45\}$ the comparison statistic was

$$t_r = \frac{\mu - \ln r}{\widehat{\mathrm{SE}}_{\mathrm{HC1}}}, \tag{45}$$

with an approximate two-sided $t_{k-1}$ reference distribution: $p = 2\Pr(T_{k-1} \geq |t_r|)$. This is the studentized comparison corresponding to the reported HC1 interval. Under this selected procedure, a reference lies outside the two-sided 95% interval exactly when its corresponding two-sided comparison gives $p < 0.05$.

These $p$-values quantify the separation from the fixed references under the weighted HC1–$t$ approximation. They inherit the approximation's small-sample and estimated-weight limitations and should not be interpreted as exactly calibrated finite-sample probabilities. They are distinct from the descriptive tail masses of individual recording bootstraps.

#### S6.6 Descriptive pooled adjacent-ratio summary

This subsection provides a descriptive summary across the two adjacent-ratio positions while retaining their position-specific analyses as the primary results.

The two adjacent-ratio positions, $r_3^2$ and $r_4^3$, were analyzed and displayed separately. A pooled adjacent-ratio summary was additionally calculated as a descriptive overview (Supplementary Fig. 4).

Repeated recordings were first combined within each animal separately for each ratio position using the procedure above. The resulting animal-by-ratio-position estimates were then entered as distinct units and assigned statistic-specific reliability weights. The pooled center was therefore recalculated from the underlying units and is not the arithmetic or geometric average of the two position-specific population centers.

Because the two ratio positions from the same animal are not independent, the pooled 95% HC1 interval is presented only as a descriptive interval under the animal-by-position unit approximation. It is not used for reference comparisons or other inferential conclusions; the position-specific animal-level analyses remain primary.

#### S6.7 Sensitivity analyses

This subsection defines the analyses used to assess dependence on the analysis unit, centering rule, weighting rule, individual units, and module-aggregation method.

We performed several sensitivity analyses, each addressing a distinct analytic choice. Their procedures are defined below; their purposes and observed implications are summarized in Supplementary Table 3, and complete numerical results are reported in Supplementary Table 6 and Section S7.

**Recordings as units.** The complete population calculation was repeated without combining recordings within animals. Each recording was treated as one analysis unit, and the same regularized center, spread, HC1, and typical-flank formulas were applied. This analysis tests whether hierarchical combination materially changes the estimated center. Because repeated recordings from the same animal are treated as separate units, narrower recording-unit intervals are not interpreted as stronger population inference.

**Bootstrap-median centering.** The raw animal estimate $y_a$ was replaced by the median $m_a$ of its MC distribution while retaining the same primary weights. The raw estimate uses the complete observed cell set, whereas the Monte Carlo median describes the center of repeated empirical resamples conditional on successful module recovery. For a smooth, well-centered estimator these quantities should be similar, but displacement can arise from nonlinearity, influential cells, finite-sample effects, or the retention rule. The bootstrap median was therefore a centering sensitivity, not an automatic bias correction.

**Equal-unit weighting.** At the animal level, the combination of repeated recordings was retained, but every animal received equal weight $1/k$. In the recording-as-unit analysis, the corresponding sensitivity assigned equal weight $1/n$ to every recording. These analyses separate the choice of independent unit from the conditional-precision weighting rule. Equal weighting deliberately ignores differences in conditional precision and was therefore treated as a robustness anchor rather than as the primary estimator.

**CNN training-seed sensitivity.** To assess sensitivity to CNN retraining stochasticity, the complete CNN analysis was repeated using 10 independently initialized networks trained with the same training data, architecture, and optimization settings. Seed 42 was the first trained realization and had been used for the primary analysis shown here before the retraining-sensitivity analysis.

Population comparisons applied the same method-level exclusions to every realization and used the common recording cohort that was retained in all 10 realizations. On this cohort, the complete primary downstream population procedure was repeated separately for each realization using its own raw estimates, bootstrap distributions, recording and animal reliability weights, and within-animal Monte Carlo combinations. Variation among the 10 resulting population centers was summarized by their geometric mean, SD in log space, and minimum–maximum range, together with the consistency of the realization-specific HC1 comparisons with unity and 49/45. The minimum lower and maximum upper HC1 limits were additionally reported as a descriptive envelope rather than as a separately calibrated confidence interval.

Separately, recording-level variation across realizations was summarized by the SD of log raw RoR and the maximum-to-minimum relative range for each recording.

**Leave-one-unit-out influence analysis.** For animal-level specifications, each animal was omitted in turn. For recording-level specifications, each recording was omitted in turn. After every omission, the median uncertainty floor, reliability weights, weighted center, HC1 standard error, degrees of freedom, and both reference comparisons were recalculated from the remaining units. The stored recording bootstraps and the combinations for the remaining animals were not regenerated. This is therefore a leave-one-unit-out population reanalysis rather than an end-to-end rerun of the upstream cell and module estimators. Primary influence and weight-concentration diagnostics are summarized in Section S7; complete results across specifications are reported in Supplementary Table 6.

##### AI declaration.

The authors used generative AI to assist with the development and debugging of analysis code, literature review, and with drafting and revising portions of the manuscript. The authors critically reviewed and edited all relevant outputs and take full responsibility for the analyses, interpretations, and content of the manuscript.

### Supplementary Discussion

#### S7 Bootstrap diagnostics, reliability, and robustness analyses

This section evaluates whether the results depend on limited module support, raw–bootstrap centering, reliability weighting, influential units, or the method used to estimate module centers.

The retained recording bootstrap contains information that the scalar reliability weight does not: its two flanks describe asymmetry, and $\pi_i$ locates the raw estimate within the retained resampling distribution. Recording-level intervals are displayed in Supplementary Fig. 4 for the adjacent ratios and in Supplementary Fig. 5b–e for RoR; the corresponding RoR diagnostics are reported numerically in Supplementary Table 5.

**Recordings with limited module support.** The two SAC recordings with the fewest retained cells, `24365_2` ($n = 42$) and `25127_1` ($n = 44$), also have the broadest RoR recording-bootstrap intervals (Supplementary Fig. 1c; Supplementary Fig. 5d; Supplementary Table 5). Both intervals contain unity and 49/45, so neither recording individually discriminates strongly between the two hypotheses. Both recordings did not enter the CNN RoR cohort because the required three-module representation was not recovered in the CNN analysis.

In `25127_1`, only one retained cell was assigned to the middle fitted module in the raw estimate, and only 3,014 of 5,000 bootstrap replicates recovered the required three-module structure. The recording remains included and visible because no independent threshold justified excluding a fitted module supported by a particular number of cells. It is the only included SAC recording from

**Supplementary Table 3.** Role of the sensitivity and validation analyses.

| Analysis | Question answered | Interpretation |
|---|---|---|
| Recordings as units | Does combining repeated recordings within an animal move the center? | Centers are nearly unchanged. Because repeated recordings are treated as independent units, the recording-unit intervals are not interpreted as stronger population inference. |
| Bootstrap-median centering | Does raw–bootstrap centering change the result? | Little effect for SAC and CNN. |
| Equal-unit weighting | Does reliability weighting determine the numerical center? | Changes the numerical center only modestly, while deliberately ignoring differences in conditional precision. |
| Leave-one-unit-out, ESS, maximum weight | Is a result dominated by one animal or recording, or by concentrated weights? | Shows that the population result is not driven by a single analysis unit, while ESS and maximum weight quantify the effective amount and concentration of independent information. |
| End-to-end synthetic benchmarks | Can the cell-scale estimation and module-aggregation pipeline recover decreasing curvature when it is present? | Validates detection under the simulated finite-arena regime, not all biological distortions. |
| KDE peak aggregation | Does the choice of GMM component centers versus KDE peak locations determine the conclusion? | KDE shifts the RoR centers downward but preserves the result against the tested 49/45 bound; the relation to unity is aggregation-dependent. |
| CNN retraining sensitivity | Does CNN retraining stochasticity materially alter recording or population RoRs? | Produces modest recording- and population-level variation but does not alter either population-level reference comparison on the fixed common cohort. |

animal `25127`, so its within-animal recording weight is $\alpha_i = 1$. Its broad propagated animal-level uncertainty yields a population weight of only $\beta_a \approx 0.0012$ in the primary SAC analysis.

Session `29502_1` illustrates a different limitation. Its largest fitted module contains only 11 SAC cells and 23 CNN cells, with fitted centers of 142.5 and 150.0 cm, respectively (Supplementary Fig. 1c,d and Supplementary Table 4). For CNN, the raw RoR is 0.9176, while the bootstrap median is 0.9222 and the central 95% interval is 0.8143–0.9613, explicitly indicating substantial uncertainty in the recording-level estimate. For SAC, the corresponding bootstrap is narrower despite the smaller component support (raw RoR: 0.839, bootstrap median: 0.832, central interval: 0.776–0.871). This narrow interval quantifies stability conditional on the observed SAC estimates; it does not remove the separate concern that SAC may become downward-biased at large grid scales (Supplementary Fig. 2a). Downward bias of the largest module center would, all else equal, reduce $r_3^2$ and consequently reduce RoR. The low SAC estimate for `29502_1` is therefore retained but interpreted cautiously rather than as definitive evidence for a comparably low biological RoR.

**Raw–bootstrap agreement.** For every included recording, the raw adjacent-ratio and RoR estimates lay within their corresponding retained 95% bootstrap intervals (Supplementary Table 5). Raw estimates were generally close to the bootstrap medians, although recording-level differences of approximately 0.01 in RoR occurred in several cases. Such differences may arise from influential cells, nonlinear GMM refitting, component instability, or conditioning on successful three-component recovery; the available analyses do not identify a single mechanism.

At the population level, bootstrap-median centering produced only small changes and did not alter the conclusion relative to unity or to the tested peak-selection reference (Supplementary Table 6).

**Relation to reliability weighting.** The reliability weights contain neither the observed RoR nor either theoretical reference and were not defined by proximity to unity or to 49/45. Nevertheless, bootstrap width and the full-data estimate are obtained from the same recording and need not be statistically independent. Weighting can therefore move the population center when recordings or animals with different RoRs also have different conditional uncertainties.

The equal-unit analyses assess dependence on the reliability-weighting rule, while the recording-as-unit and leave-one-unit-out analyses assess dependence on the hierarchical unit and individual units, respectively. The estimated reliability weights were treated as fixed after construction; uncertainty in the weights themselves was not propagated.

**Sensitivity to CNN retraining.** Across the eight-recordings cohort, the SD of log raw RoR across CNN realizations corresponded to approximately 0.6–1.9% one-SD proportional variation, while the observed maximum-to-minimum relative ranges were 1.8 to 6.1%. The largest one-SD proportional variation occurred for recording `29502_1` (1.9%), and the largest relative range also occurred for recording `29502_1` (6.1%).

The 10 complete population reruns yielded RoR centers from 0.9674 to 0.9926, with an across-realization geometric mean of 0.9843 and an SD of log population RoR of 0.0075 (Supplementary Table 7). Every realization-specific HC1 interval included unity and excluded 49/45; collectively, the descriptive envelope of the interval limits spanned 0.9216–1.0391. Thus, CNN retraining produced modest recording- and population-level variation but did not alter either reference comparison.

**Sensitivity to KDE-based module aggregation.** In a matched-cohort sensitivity analysis, replacing the GMM component centers with KDE peak locations shifted the estimated RoR downward, particularly for SAC. In the primary reliability-weighted animal analysis, the KDE population centers were 0.9449 for SAC (approximate HC1 95% CI, 0.9094–0.9818; $p = 0.010$ versus unity) and 0.9871 for CNN (0.9657–1.0090; $p = 0.177$ versus unity). Both intervals lay entirely below the tested peak-selection reference 49/45 ($p = 5.09 \times 10^{-5}$ and $p = 2.42 \times 10^{-4}$ for SAC and CNN, respectively), and this conclusion persisted across the weighting, centering, analysis-unit, and leave-one-unit-out specifications.

The relation to unity was less invariant. The primary GMM intervals included unity for both estimators. Under KDE aggregation, the primary SAC interval excluded unity, whereas the CNN interval continued to include it; both equal-weight KDE sensitivity for SAC also included unity. Thus, the conclusion relative to the tested peak-selection reference is robust to GMM versus KDE aggregation, whereas the precise relation to unity, particularly for SAC, depends on the aggregation and weighting specifications. In the synthetic three-module benchmarks, the CNN-based pipelines recovered the known RoR more closely than the corresponding SAC-based pipelines under both the constant- and decreasing-ratio conditions (Supplementary Fig. 3b,c). Where the empirical estimators differ, these benchmark results lend greater weight to the CNN-based inference.

**Sensitivity and influence summary.** Bootstrap-median centering yielded an RoR center of 0.9603 for SAC (approximate HC1 95% interval, 0.9104–1.0129) and 0.9958 for CNN (0.9717–1.0205). Under equal-animal weighting, the CNN center was 0.9767; the complete equal-weight and recording-as-unit results are reported in Supplementary Table 6. Within the primary reliability-weighted animal analysis, the leave-one-animal-out population centers ranged from 0.9554 to 0.9811 for SAC and from 0.9830 to 0.9941 for CNN. Complete leave-one-unit-out ranges for all reported specifications are provided in Supplementary Table 6. In the full primary analyses, the effective sample sizes were 5.59 of 8 animals for SAC and 3.74 of 5 animals for CNN, and the corresponding maximum animal weights were 0.220 and 0.328. Together with the KDE analysis, these diagnostics quantify sensitivity to the analysis unit, centering rule, weighting rule, individual animals, weight concentration, and module-aggregation method. Their implications for the theoretical references are summarized in Section S8.

#### S8 Interpretation and limits of the ratio-of-ratios result

This section distinguishes the robust conclusion relative to the tested peak-selection bound from the somewhat less stable relation to unity and states the principal estimator-specific and population-level limitations.

**The robust empirical conclusion.** Every included recording has a raw RoR below the tested peak-selection lower bound $49/45 \approx 1.089$ (maximum $\approx 1.044$; Supplementary Fig. 5d,e and Supplementary Table 4). The recording-bootstrap interval lies fully below this bound in all 8 CNN recordings and in 9 of 11 SAC recordings (Supplementary Fig. 5d,e and Supplementary Table 5), independently of any population weighting. The primary animal-level HC1 intervals also exclude 49/45. The full-data recording-as-unit, equal-animal-weight, equal-recording-weight, and bootstrap-median specifications reported in Supplementary Table 6 all retain population centers below this reference, with approximate HC1 comparison $p$-values below $p = 0.05$. Therefore, the observed module triples provide strong evidence against the tested 49/45 lower bound under the theory-conditional $(M_2, M_3, M_4)$ indexing.

The relation to unity is less invariant. The primary reliability-weighted population centers are 0.963 for the SAC analysis (approximate HC1 95% CI, 0.914–1.015) and 0.990 for the CNN analysis (0.969–1.012). Both intervals contain RoR $= 1$ and lie entirely below the conservative peak-selection bound $49/45 \approx 1.089$. For SAC, the reliability-weighted recording-as-unit specification narrowly excluded unity ($p = 0.045$), whereas the animal-level and equal-weight specifications did not. CNN training-seed sensitivity is considered separately under “Sensitivity to CNN retraining” in Section S7.

The end-to-end synthetic benchmarks recovered distinct locally constant- and unequal-ratio structures, showing that the pipeline does not mechanically force estimates toward unity (Sections S3 and S5; Supplementary Fig. 3b,c). Across the sensitivity analyses, none of the tested analysis-unit, weighting, or centering specifications changed the conclusion relative to 49/45 (Section S7; Supplementary Table 6). Reliability weighting is retained as the primary specification because it accounts for the marked differences in conditional precision, with equal weighting serving as a robustness analysis.

**Relation to geometric-progression accounts.** A constant adjacent ratio, and hence RoR $= 1$, is a property shared by theories that generate a geometric sequence of grid scales[8,9]. The measured adjacent-ratio centers of approximately 1.4–1.5, together with RoR values near one, are compatible with that class of local scale organization. They do not uniquely establish optimization of decoding accuracy as the biological mechanism, because other developmental or network processes may also generate approximately geometric module centers.

Conversely, the present result provides strong evidence against theories that predict an RoR at or above 49/45.

**Estimator-specific limitations.** SAC and CNN do not provide numerically identical evidence relative to unity. The SAC analysis yields a lower population center and a stronger difference between its two adjacent-ratio estimates, but synthetic validation identifies a scale-dependent downward bias at the largest scales. Such bias would tend to lower $r_3^2$ and RoR (Supplementary Fig. 2). The CNN analysis substantially reduces this ratio asymmetry under the primary full-data specification.

**A population center does not imply per-animal exactness.** The reported population values are reliability-weighted centers of 5 CNN or 8 SAC animal estimates. Individual animal estimates vary, and the observed spread combines biological heterogeneity with remaining measurement error. With the present number of animals, the analysis cannot estimate latent between-animal heterogeneity precisely, determine how broadly the estimates generalize beyond these datasets, or identify the biological mechanism producing the observed scale relation. The present analysis should therefore be read as a quantitative empirical constraint from the highest-density suitable recordings analyzed here, conditional on the stated estimators, module assignment, and aggregation procedure.

### Supplementary Tables

**Supplementary Table 4.** Included recordings, selected GMM module centers, adjacent ratios, and raw ratio-of-ratios. Module centers are shown in cm and ordered from largest to smallest for each recording.

| Dataset | Method | Recording | $M_2$ | $M_3$ | $M_4$ | $r_3^2$ | $r_4^3$ | $\mathrm{RoR}_4^2$ |
|---|---|---|---|---|---|---|---|---|
| Gardner | CNN | R_d1 | 127.5 | 83.9 | 57.2 | 1.5188 | 1.4677 | 1.0348 |
| Gardner | CNN | R_d2 | 112.4 | 78.9 | 53.4 | 1.4238 | 1.4771 | 0.9639 |
| Vollan | CNN | 25843_1 | 127.8 | 84.3 | 57.2 | 1.5164 | 1.4726 | 1.0297 |
| Vollan | CNN | 25843_2 | 114.5 | 79.3 | 53.8 | 1.4444 | 1.4737 | 0.9801 |
| Vollan | CNN | 28258_4 | 118.7 | 78.3 | 51.0 | 1.5161 | 1.5349 | 0.9877 |
| Vollan | CNN | 28304_1 | 143.2 | 97.8 | 64.9 | 1.4638 | 1.5074 | 0.9711 |
| Vollan | CNN | 28304_2 | 137.6 | 93.1 | 61.7 | 1.4788 | 1.5071 | 0.9812 |
| Vollan | CNN | 29502_1 | 150.0 | 106.5 | 69.4 | 1.4081 | 1.5346 | 0.9176 |
| Gardner | SAC | R_d1 | 123.3 | 84.3 | 57.4 | 1.4622 | 1.4693 | 0.9952 |
| Gardner | SAC | R_d2 | 113.3 | 78.6 | 53.9 | 1.4407 | 1.4586 | 0.9877 |
| Vollan | SAC | 24365_2 | 90.9 | 60.8 | 42.2 | 1.4949 | 1.4388 | 1.0390 |
| Vollan | SAC | 25127_1 | 120.6 | 71.6 | 44.4 | 1.6833 | 1.6131 | 1.0435 |
| Vollan | SAC | 25843_1 | 124.5 | 84.5 | 57.5 | 1.4732 | 1.4701 | 1.0022 |
| Vollan | SAC | 25843_2 | 113.4 | 80.5 | 54.3 | 1.4080 | 1.4834 | 0.9492 |
| Vollan | SAC | 26018_2 | 89.3 | 64.5 | 46.9 | 1.3832 | 1.3773 | 1.0043 |
| Vollan | SAC | 28258_4 | 115.1 | 76.5 | 51.7 | 1.5051 | 1.4798 | 1.0171 |
| Vollan | SAC | 28304_1 | 139.8 | 99.5 | 64.7 | 1.4056 | 1.5377 | 0.9141 |
| Vollan | SAC | 28304_2 | 134.1 | 93.2 | 62.0 | 1.4385 | 1.5042 | 0.9563 |
| Vollan | SAC | 29502_1 | 142.5 | 108.3 | 69.0 | 1.3161 | 1.5683 | 0.8392 |

**Supplementary Table 5.** Recording-level cell-bootstrap audit and primary RoR reliability quantities. Raw is the full-data estimate. Bootstrap median and bootstrap 95% interval summarize the retained cell-bootstrap replicates conditional on recovery of the required three-module structure. Bootstrap retained/requested gives the number of usable bootstrap replicates among the requested draws, and raw bootstrap percentile is the percentage of retained bootstrap values less than or equal to the raw estimate. The final two columns are reported only for RoR and refer to the primary reliability-weighted animal analysis. In each stacked cell, the upper value is the unregularized longer flank ($h_i$ at recording level or $H_a$ after within-animal Monte Carlo combination), and the lower value is the corresponding regularized normalized weight ($\alpha_i$ within animal or $\beta_a$ across animals). The method-specific median-flank floors enter the weights but not the displayed flanks. $H_a$ and $\beta_a$ are repeated for recordings belonging to the same animal; $\alpha_i = 1$ for an animal represented by one included recording and does not imply maximal population reliability.

| Method | Recording | Statistic | Raw | Recording bootstrap | | | | Primary RoR reliability | |
|---|---|---|---|---|---|---|---|---|---|
| | | | | Median | 95% interval | Retained/<br>requested | Raw<br>percentile (%) | $h_i$<br>$\alpha_i$ | $H_a$<br>$\beta_a$ |
| CNN | R_d1 | $r_3^2$ | 1.5188 | 1.5164 | 1.4840–1.5484 | 4379/5000 | 55.8 | – | – |
| | | $r_4^3$ | 1.4677 | 1.4683 | 1.4554–1.4807 | 4379/5000 | 46.8 | – | – |
| | | $\mathrm{RoR}_4^2$ | 1.0348 | 1.0330 | 1.0050–1.0605 | 4379/5000 | 55.2 | 0.0275<br>0.565 | 0.0222<br>0.328 |
| CNN | R_d2 | $r_3^2$ | 1.4238 | 1.4424 | 1.3894–1.4755 | 5000/5000 | 16.7 | – | – |
| | | $r_4^3$ | 1.4771 | 1.4766 | 1.4654–1.4868 | 5000/5000 | 54.2 | – | – |
| | | $\mathrm{RoR}_4^2$ | 0.9639 | 0.9769 | 0.9404–1.0026 | 5000/5000 | 18.0 | 0.0381<br>0.435 | 0.0222<br>0.328 |
| CNN | 25843_1 | $r_3^2$ | 1.5164 | 1.5323 | 1.4623–1.5690 | 4289/5000 | 22.9 | – | – |
| | | $r_4^3$ | 1.4726 | 1.4702 | 1.4543–1.4834 | 4289/5000 | 62.2 | – | – |
| | | $\mathrm{RoR}_4^2$ | 1.0297 | 1.0426 | 0.9889–1.0741 | 4289/5000 | 23.8 | 0.0529<br>0.430 | 0.0328<br>0.239 |
| CNN | 25843_2 | $r_3^2$ | 1.4444 | 1.4562 | 1.3984–1.4930 | 4983/5000 | 29.7 | – | – |
| | | $r_4^3$ | 1.4737 | 1.4732 | 1.4632–1.4815 | 4983/5000 | 54.5 | – | – |
| | | $\mathrm{RoR}_4^2$ | 0.9801 | 0.9884 | 0.9482–1.0165 | 4983/5000 | 30.3 | 0.0415<br>0.570 | 0.0328<br>0.239 |
| CNN | 28258_4 | $r_3^2$ | 1.5161 | 1.5354 | 1.4455–1.6278 | 3732/5000 | 31.9 | – | – |
| | | $r_4^3$ | 1.5349 | 1.5363 | 1.5062–1.5834 | 3732/5000 | 46.4 | – | – |
| | | $\mathrm{RoR}_4^2$ | 0.9877 | 0.9989 | 0.9375–1.0612 | 3732/5000 | 34.8 | 0.0634<br>1.000 | 0.0635<br>0.100 |
| CNN | 28304_1 | $r_3^2$ | 1.4638 | 1.4660 | 1.4198–1.5061 | 5000/5000 | 45.9 | – | – |
| | | $r_4^3$ | 1.5074 | 1.5076 | 1.4898–1.5268 | 5000/5000 | 48.8 | – | – |
| | | $\mathrm{RoR}_4^2$ | 0.9711 | 0.9721 | 0.9392–1.0026 | 5000/5000 | 47.7 | 0.0345<br>0.507 | 0.0248<br>0.303 |
| CNN | 28304_2 | $r_3^2$ | 1.4788 | 1.4845 | 1.4365–1.5307 | 5000/5000 | 42.2 | – | – |
| | | $r_4^3$ | 1.5071 | 1.5079 | 1.4912–1.5262 | 5000/5000 | 46.7 | – | – |
| | | $\mathrm{RoR}_4^2$ | 0.9812 | 0.9848 | 0.9504–1.0183 | 5000/5000 | 42.5 | 0.0356<br>0.493 | 0.0248<br>0.303 |
| CNN | 29502_1 | $r_3^2$ | 1.4081 | 1.4216 | 1.2418–1.4705 | 4844/5000 | 35.9 | – | – |
| | | $r_4^3$ | 1.5346 | 1.5366 | 1.5098–1.5710 | 4844/5000 | 44.5 | – | – |
| | | $\mathrm{RoR}_4^2$ | 0.9176 | 0.9222 | 0.8143–0.9613 | 4844/5000 | 42.3 | 0.1244<br>1.000 | 0.1263<br>0.030 |
| SAC | R_d1 | $r_3^2$ | 1.4622 | 1.4599 | 1.4336–1.4886 | 4965/5000 | 56.9 | – | – |
| | | $r_4^3$ | 1.4693 | 1.4697 | 1.4521–1.4865 | 4965/5000 | 48.0 | – | – |
| | | $\mathrm{RoR}_4^2$ | 0.9952 | 0.9932 | 0.9679–1.0221 | 4965/5000 | 55.9 | 0.0286<br>0.565 | 0.0234<br>0.220 |
| SAC | R_d2 | $r_3^2$ | 1.4407 | 1.4281 | 1.3875–1.4683 | 4764/5000 | 72.6 | – | – |
| | | $r_4^3$ | 1.4586 | 1.4618 | 1.4410–1.4796 | 4764/5000 | 37.0 | – | – |
| | | $\mathrm{RoR}_4^2$ | 0.9877 | 0.9764 | 0.9433–1.0159 | 4764/5000 | 72.0 | 0.0397<br>0.435 | 0.0234<br>0.220 |
| SAC | 24365_2 | $r_3^2$ | 1.4949 | 1.4995 | 1.3896–1.6061 | 3921/5000 | 46.0 | – | – |
| | | $r_4^3$ | 1.4388 | 1.4435 | 1.3517–1.5776 | 3921/5000 | 46.3 | – | – |

Supplementary Table 5 continued

| Method | Recording | Statistic | Raw | Recording bootstrap | | | | Primary RoR reliability | |
|---|---|---|---|---|---|---|---|---|---|
| | | | | Median | 95% interval | Retained/ requested | Raw percentile (%) | $h_i$ <br> $\alpha_i$ | $H_a$ <br> $\beta_a$ |
| | | $\mathrm{RoR}_4^2$ | 1.0390 | 1.0385 | 0.9119–1.1589 | 3921/5000 | 50.4 | 0.1300 | 0.1294 |
| | | | | | | | | 1.000 | 0.053 |
| SAC | 25127_1 | $r_3^2$ | 1.6833 | 1.6644 | 1.0296–1.7394 | 3014/5000 | 64.9 | – | – |
| | | $r_4^3$ | 1.6131 | 1.6191 | 1.1094–2.6763 | 3014/5000 | 39.7 | – | – |
| | | $\mathrm{RoR}_4^2$ | 1.0435 | 1.0331 | 0.3854–1.4874 | 3014/5000 | 61.3 | 0.9859 | 0.9794 |
| | | | | | | | | 1.000 | 0.001 |
| SAC | 25843_1 | $r_3^2$ | 1.4732 | 1.4669 | 1.4387–1.4982 | 4950/5000 | 64.8 | – | – |
| | | $r_4^3$ | 1.4701 | 1.4712 | 1.4538–1.4875 | 4950/5000 | 44.5 | – | – |
| | | $\mathrm{RoR}_4^2$ | 1.0022 | 0.9972 | 0.9707–1.0271 | 4950/5000 | 63.5 | 0.0296 | 0.0232 |
| | | | | | | | | 0.568 | 0.220 |
| SAC | 25843_2 | $r_3^2$ | 1.4080 | 1.4102 | 1.3769–1.4519 | 4770/5000 | 44.8 | – | – |
| | | $r_4^3$ | 1.4834 | 1.4823 | 1.4610–1.5021 | 4770/5000 | 53.8 | – | – |
| | | $\mathrm{RoR}_4^2$ | 0.9492 | 0.9508 | 0.9251–0.9906 | 4770/5000 | 45.4 | 0.0410 | 0.0232 |
| | | | | | | | | 0.432 | 0.220 |
| SAC | 26018_2 | $r_3^2$ | 1.3832 | 1.3861 | 1.2995–1.4321 | 4872/5000 | 45.2 | – | – |
| | | $r_4^3$ | 1.3773 | 1.3862 | 1.3527–1.5052 | 4872/5000 | 32.3 | – | – |
| | | $\mathrm{RoR}_4^2$ | 1.0043 | 0.9997 | 0.8636–1.0495 | 4872/5000 | 58.0 | 0.1463 | 0.1478 |
| | | | | | | | | 1.000 | 0.043 |
| SAC | 28258_4 | $r_3^2$ | 1.5051 | 1.5045 | 1.4300–1.5729 | 3881/5000 | 50.5 | – | – |
| | | $r_4^3$ | 1.4798 | 1.4807 | 1.4417–1.5166 | 3881/5000 | 48.0 | – | – |
| | | $\mathrm{RoR}_4^2$ | 1.0171 | 1.0158 | 0.9515–1.0816 | 3881/5000 | 51.6 | 0.0655 | 0.0659 |
| | | | | | | | | 1.000 | 0.127 |
| SAC | 28304_1 | $r_3^2$ | 1.4056 | 1.4038 | 1.3602–1.4459 | 4620/5000 | 53.6 | – | – |
| | | $r_4^3$ | 1.5377 | 1.5373 | 1.5075–1.5610 | 4620/5000 | 51.2 | – | – |
| | | $\mathrm{RoR}_4^2$ | 0.9141 | 0.9134 | 0.8809–0.9473 | 4620/5000 | 51.4 | 0.0364 | 0.0249 |
| | | | | | | | | 0.486 | 0.217 |
| SAC | 28304_2 | $r_3^2$ | 1.4385 | 1.4373 | 1.3995–1.4737 | 4993/5000 | 52.7 | – | – |
| | | $r_4^3$ | 1.5042 | 1.5033 | 1.4790–1.5276 | 4993/5000 | 52.9 | – | – |
| | | $\mathrm{RoR}_4^2$ | 0.9563 | 0.9559 | 0.9240–0.9891 | 4993/5000 | 51.1 | 0.0341 | 0.0249 |
| | | | | | | | | 0.514 | 0.217 |
| SAC | 29502_1 | $r_3^2$ | 1.3161 | 1.3035 | 1.2123–1.3555 | 4608/5000 | 65.6 | – | – |
| | | $r_4^3$ | 1.5683 | 1.5630 | 1.5366–1.5917 | 4608/5000 | 64.6 | – | – |
| | | $\mathrm{RoR}_4^2$ | 0.8392 | 0.8321 | 0.7761–0.8707 | 4608/5000 | 63.4 | 0.0697 | 0.0700 |
| | | | | | | | | 1.000 | 0.119 |

**Supplementary Table 6.** RoR population sensitivity and leave-one-unit-out analyses. Centers, HC1 intervals, and reference comparisons are calculated in log space and back-transformed for display. Reliability-weighted rows use the median longer flank as the regularization floor; the bootstrap-median rows change the animal point values while retaining adaptive weighting; and equal-weight rows assign weight $1/k$ to every unit. Each indented leave-one-out subrow gives the range obtained after omitting one analysis unit at a time—one animal for animal specifications and one recording for recording specifications. For adaptive specifications, the floor and normalized weights are recalculated after each omission, together with the HC1 standard error, degrees of freedom, and reference comparisons.

| Method | Specification | Center | HC1 95% CI | $p$ vs 1 | $p$ vs 49/45 | ESS / max weight |
|---|---|---|---|---|---|---|
| CNN | **Reliability-weighted animals** | 0.99022 | 0.9688–1.0121 | 0.2806 | 0.000272 | 3.74 / 0.328 |
| | Leave one animal out: center 0.9830–0.9941<br>$p$ vs 1: 0.1715–0.5087; $p$ vs 49/45: 0.001331–0.002365 | | | | | |
| CNN | **Reliability-weighted recordings** | 0.99049 | 0.9634–1.0183 | 0.4417 | $8.544 \times 10^{-5}$ | 6.91 / 0.191 |
| | Leave one recording out: center 0.9803–0.9950<br>$p$ vs 1: 0.0515–0.7019; $p$ vs 49/45: $1.385 \times 10^{-5}$–0.000503 | | | | | |
| CNN | **Bootstrap-median animals** | 0.99580 | 0.9717–1.0205 | 0.6581 | 0.000534 | 3.74 / 0.328 |
| | Leave one animal out: center 0.9894–1.0014<br>$p$ vs 1: 0.4218–0.8646; $p$ vs 49/45: 0.001638–0.003608 | | | | | |
| CNN | **Equal-weight animals** | 0.97665 | 0.9333–1.0221 | 0.2224 | 0.002661 | 5.00 / 0.200 |
| | Leave one animal out: center 0.9701–0.9920<br>$p$ vs 1: 0.2131–0.3475; $p$ vs 49/45: 0.000705–0.0143 | | | | | |
| CNN | **Equal-weight recordings** | 0.98265 | 0.9519–1.0144 | 0.2337 | 0.000122 | 8.00 / 0.125 |
| | Leave one recording out: center 0.9754–0.9923<br>$p$ vs 1: 0.1031–0.4949; $p$ vs 49/45: 0.000124–0.0006 | | | | | |
| SAC | **Reliability-weighted animals** | 0.96327 | 0.9140–1.0152 | 0.136 | 0.000891 | 5.59 / 0.220 |
| | Leave one animal out: center 0.9554–0.9811<br>$p$ vs 1: 0.1118–0.3189; $p$ vs 49/45: 0.000397–0.005497 | | | | | |
| SAC | **Reliability-weighted recordings** | 0.96553 | 0.9331–0.9990 | 0.045 | $1.387 \times 10^{-5}$ | 7.94 / 0.160 |
| | Leave one recording out: center 0.9586–0.9744<br>$p$ vs 1: 0.0378–0.1146; $p$ vs 49/45: $1.264 \times 10^{-5}$–0.000116 | | | | | |
| SAC | **Bootstrap-median animals** | 0.96029 | 0.9104–1.0129 | 0.1154 | 0.000839 | 5.59 / 0.220 |
| | Leave one animal out: center 0.9524–0.9788<br>$p$ vs 1: 0.0956–0.2777; $p$ vs 49/45: 0.000289–0.005379 | | | | | |
| SAC | **Equal-weight animals** | 0.97906 | 0.9223–1.0393 | 0.4296 | 0.003981 | 8.00 / 0.125 |
| | Leave one animal out: center 0.9702–1.0009<br>$p$ vs 1: 0.3083–0.9537; $p$ vs 49/45: 0.001041–0.0122 | | | | | |
| SAC | **Equal-weight recordings** | 0.97532 | 0.9345–1.0179 | 0.2215 | 0.000186 | 11.00 / 0.091 |
| | Leave one recording out: center 0.9687–0.9901<br>$p$ vs 1: 0.1438–0.4681; $p$ vs 49/45: $4.894 \times 10^{-5}$–0.000627 | | | | | |

**Supplementary Table 7.** CNN training-seed sensitivity. Population estimates were recalculated for 10 independently trained CNN realizations on the same 8-recording, 5-animal cohort. Each realization was analyzed using its own raw estimates, bootstrap distributions, recording weights, within-animal Monte Carlo combinations, and animal weights. Seed 42 was the first trained realization and had been used for the primary analysis before the retraining-sensitivity analysis. The final row reports the geometric mean and minimum–maximum range of the realization-specific population centers, the descriptive envelope of the HC1 limits, and the ranges of the two-sided reference-comparison $p$-values.

| CNN realization | Population RoR | HC1 95% interval | $p$ vs. unity | $p$ vs. 49/45 |
|---|---|---|---|---|
| Seed 42 (primary) | 0.9902 | 0.9688–1.0121 | 0.2806 | 0.0003 |
| Seed 43 | 0.9876 | 0.9594–1.0165 | 0.2950 | 0.0007 |
| Seed 44 | 0.9884 | 0.9418–1.0374 | 0.5408 | 0.0051 |
| Seed 45 | 0.9865 | 0.9566–1.0174 | 0.2894 | 0.0009 |
| Seed 46 | 0.9926 | 0.9686–1.0172 | 0.4471 | 0.0005 |
| Seed 47 | 0.9872 | 0.9379–1.0391 | 0.5246 | 0.0060 |
| Seed 48 | 0.9674 | 0.9216–1.0155 | 0.1305 | 0.0025 |
| Seed 49 | 0.9828 | 0.9380–1.0298 | 0.3614 | 0.0037 |
| Seed 50 | 0.9832 | 0.9457–1.0223 | 0.2948 | 0.0019 |
| Seed 51 | 0.9768 | 0.9436–1.0111 | 0.1323 | 0.0009 |
| Across realizations | Geometric mean 0.9843<br>range 0.9674–0.9926 | Envelope<br>0.9216–1.0391 | Range<br>0.1305–0.5408 | Range<br>0.0003–0.0060 |

### Supplementary Figures

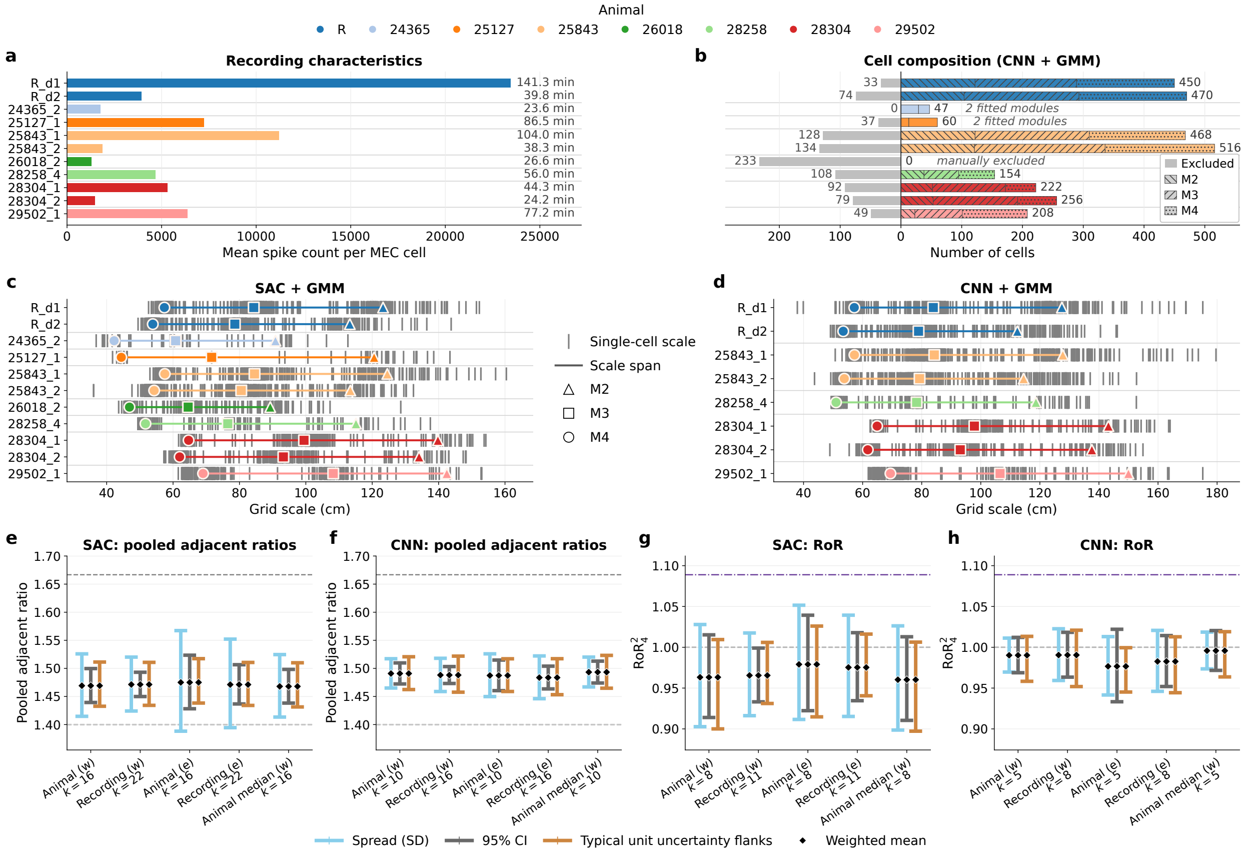


**Supplementary Figure 1. Recording-level scale structure and population-summary sensitivity analyses.** (**a**) Mean spike count per MEC cell for each recording. Bar color identifies the animal, and the recording duration is printed at the right. (**b**) CNN+GMM cell composition for the same recording set. Excluded cells are shown to the left of zero in gray; retained cells assigned to modules M2–M4 are stacked to the right, with overlaid patterns corresponding to identified modules. The number at the right gives the retained-cell count, and italic text reports the fitted module count when the recording did not meet the three-module inclusion criterion. (**c**,**d**) Single-cell grid scales and the three fitted GMM module representatives for the SAC and CNN analyses, respectively. Gray vertical ticks show individual retained-cell scales, the horizontal segment spans the selected module centers, and the triangle, square, and circle denote M2, M3, and M4, respectively. Symbol color identifies the animal. Recordings with limited module support remain included and visible. Module-specific CNN support is shown in panel **b**; total retained-cell counts, fitted centers, raw ratios, and recording-bootstrap diagnostics are reported in Supplementary Tables 4 and 5 and discussed in Section S7. (**e**,**f**) SAC and CNN population summaries of the pooled adjacent ratios under five specifications: reliability-weighted animals, reliability-weighted recordings, equal-weight animals, equal-weight recordings, and reliability-weighted animals after replacing each raw animal estimate by its Monte Carlo median. Blue intervals show the weighted observed-unit spread, dark-gray intervals the approximate descriptive 95% HC1 interval for the center, and orange-brown intervals show the typical conditional-uncertainty flanks of the chosen analysis unit; diamonds mark the weighted geometric center. Gray dashed lines delimit the peak-selection range $7/5 \le r_3^2 \le 5/3$ over $f \in [-\frac{1}{2}, \frac{1}{2}]$.[11] (**g**,**h**) Corresponding SAC and CNN summaries for $\mathrm{RoR}_4^2$. Here the dark-gray intervals are the approximate animal- or recording-level 95% HC1 confidence intervals defined for the corresponding specification.

The gray dashed lines mark RoR $= 1$, and the purple dash-dotted lines mark the conservative peak-selection lower bound $49/45$, with $f = 1/2$. All population calculations are performed in log space and exponentiated for display; the spread and interval limits therefore represent multiplicative deviations around geometric means.

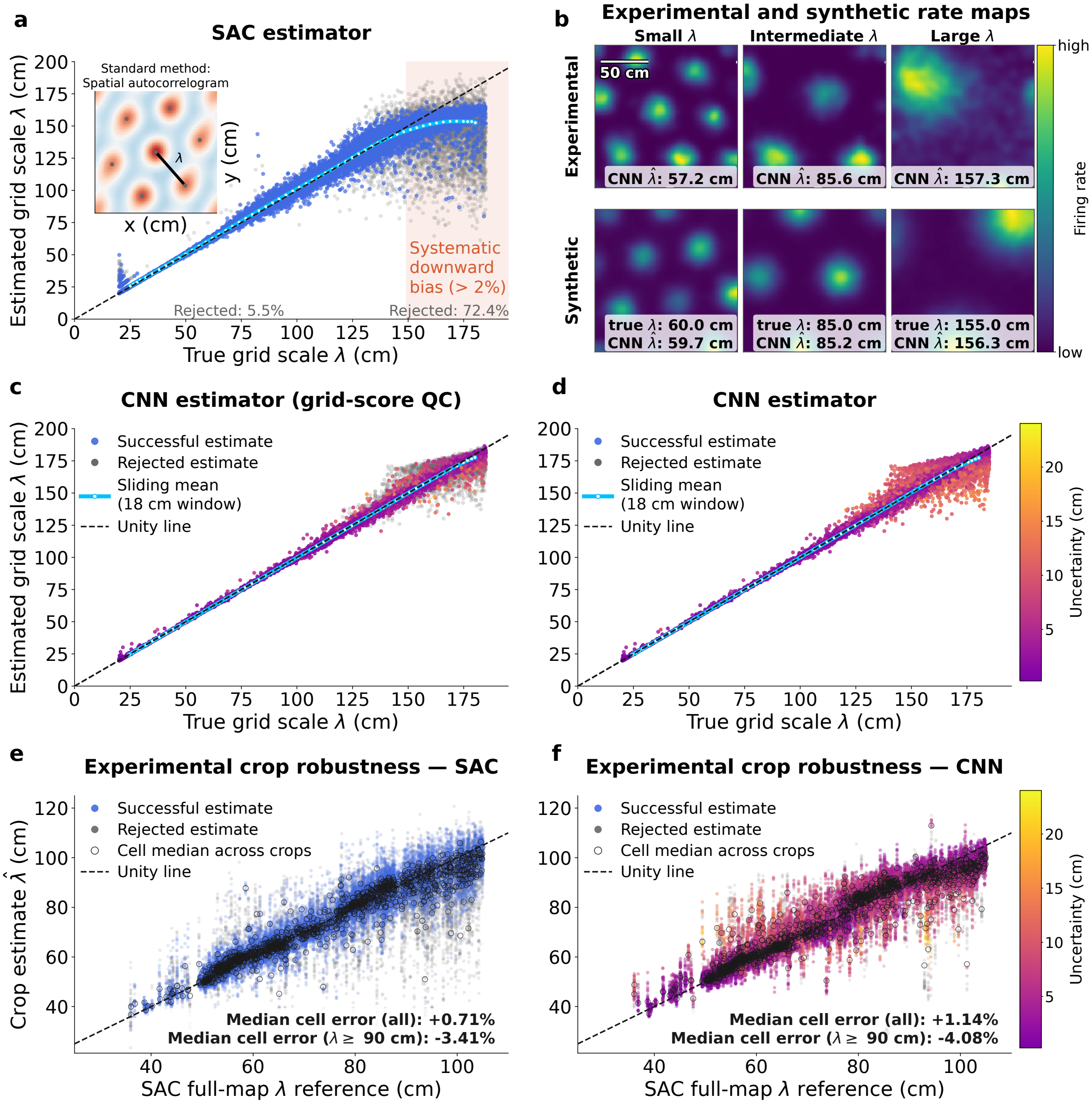


**Supplementary Figure 2. Validation and comparison of single-cell grid-scale estimators.** (**a**) Performance of the spatial-autocorrelogram (SAC) estimator on simulated grid-cell rate maps. Each point represents one simulated cell and shows the estimated grid scale as a function of its ground-truth scale. Blue and gray points denote estimates that passed or failed the SAC quality-control criteria, respectively. The cyan line shows the sliding-window mean estimate, and the black dashed line marks equality between estimated and true grid scale. The shaded region indicates the range in which the mean estimate shows a systematic downward bias exceeding 2%. The inset illustrates the SAC procedure using a spatial autocorrelogram; marked peaks indicate the first ring of autocorrelation maxima, and the radial segment denotes the inferred grid scale $\lambda$. (**b**) Example experimental and simulated rate maps used for estimator evaluation. The upper panels show three experimental grid-cell rate maps ordered by estimated grid scale, and the lower panels show three simulated examples ordered by ground-truth grid scale. Insets report the corresponding CNN estimate and, for simulated cells, the known true grid scale. (**c**) Performance of the convolutional neural-network (CNN) estimator when grid-score quality-control rejection is retained. Successful CNN estimates are colored by their estimated uncertainty, whereas cells failing the grid-score quality-control criterion are shown in gray. The cyan line shows the uncertainty-weighted sliding-window mean, and the black dashed line denotes equality between estimated and true grid scale. (**d**) CNN estimates for the same simulated cells without marking grid-score quality-control failures as rejected. All finite CNN predictions contribute to the displayed scatter and uncertainty-weighted sliding-window mean. Point color denotes the estimated uncertainty, using the color scale at the right. Comparison with panel **c** isolates the effect of the grid-score quality-control rejection criterion while keeping the CNN predictions and simulated-cell population fixed. For visual clarity, at most 30,000 simulated cells are displayed in each simulated-data scatter panel. Sliding-window summaries, uncertainty normalization, rejection fractions, bias estimates, and axis limits for the simulated panels are calculated from the complete set of 60,000 held-out simulated cells. (**e**,**f**) Experimental crop robustness of the SAC and CNN estimators, respectively. We generated 172,183 crop observations from 1,423 experimental cells, with 121 crops per cell. The horizontal axis shows the reference grid scale estimated from the corresponding complete experimental rate map using the SAC estimator. The vertical axis shows the estimate obtained from the cropped map. For visualization, 28 crops were randomly selected within each original cell, yielding 39,844 displayed observations per panel. The identical crop IDs and identical axis limits are used in panels **e** and **f**. In panel **e**, SAC estimates passing the complete SAC quality-control criteria are shown in blue and rejected estimates in gray. In panel **f**, CNN estimates passing the grid-score criterion are colored by their predicted uncertainty, using the color scale at the right, whereas grid-score failures are shown in gray. Larger open circles show the median of the crop estimates passing the corresponding method-specific quality-control criterion for each cell across its 121 crop observations, and the black dashed line marks equality between the crop estimate and the full-map reference. The two-line annotation in each panel reports the median signed cell error for all 1,423 cells and separately for the 294 cells whose SAC full-map reference scale is at least 90 cm. For each cell and method, $\Delta \log \lambda = \log(\hat{\lambda}_{\text{crop}}/\lambda_{\text{full}})$ is calculated for each crop passing the corresponding method-specific quality-control criterion and its median is taken; the displayed value is then the median across cells, transformed to a multiplicative percentage as $100[\exp(\text{median}(\Delta \log \lambda))-1]$. Positive values denote overestimation and negative values denote downward displacement relative to the full-map reference. The CNN shows a slightly larger positive signed displacement overall and a slightly larger downward displacement in the predefined large-scale subset.

Across all experimental crops, 135,299 SAC estimates (78.6%) passed SAC quality control, whereas 149,373 crops (86.8%) passed the CNN panel's grid-score criterion.

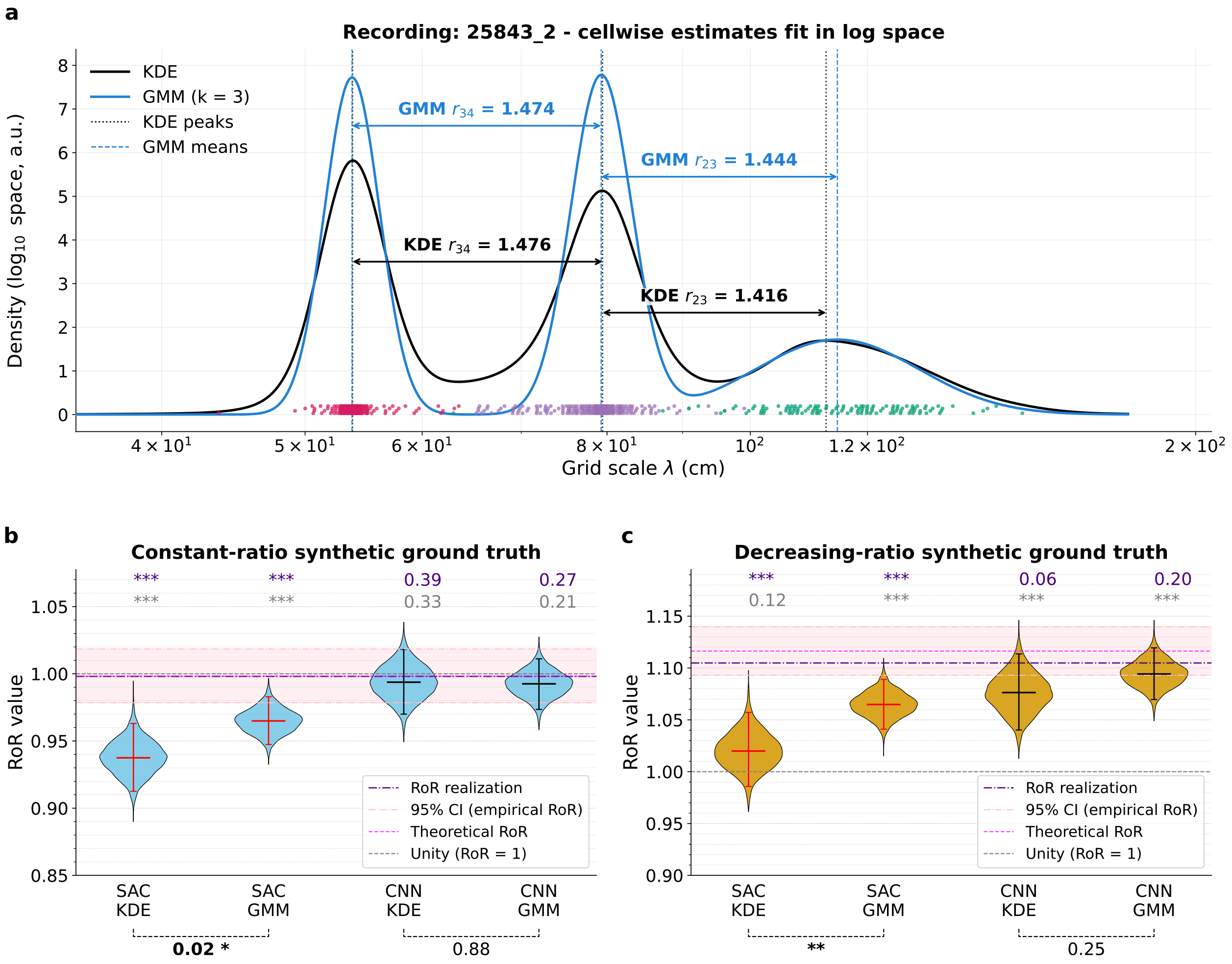


**Supplementary Figure 3. Module-assignment overview and synthetic pipeline validation.** (**a**) Example extraction of consecutive grid-scale ratios from the cellwise grid-scale estimates of a representative Vollan recording (25843_2), obtained using the CNN-based scale estimator. The distributions were fitted in log space, while the horizontal axis shows grid scales $\lambda$ in centimeters on a logarithmic scale. The black solid curve is the kernel-density estimate (KDE), and the blue solid curve is the fitted three-component Gaussian mixture model (GMM). Black dotted and blue dashed vertical lines mark the KDE peak positions and GMM component means, respectively. Colored rug markers near the baseline show the retained cellwise grid-scale estimates, with colors indicating their assigned grid modules. Horizontal arrows show the consecutive scale ratios $r_4^3$ and $r_3^2$ obtained from the KDE peaks (black) or GMM means (blue). (**b**,**c**) Bootstrap distributions of the $\mathrm{RoR}_4^2$ values obtained by applying four pipelines (SAC+KDE, SAC+GMM, CNN+KDE, and CNN+GMM) to the synthetic three-module benchmarks: (**b**) the constant-ratio condition and (**c**) the decreasing-ratio condition. The violins show the retained bootstrap distributions (5,000 resamples per violin); the central horizontal line and whiskers indicate the bootstrap median and the central 95% percentile bootstrap confidence interval, respectively. The gray dashed line denotes unity, RoR = 1, whereas the purple dash-dotted line denotes the RoR realization in the generated benchmark dataset. These are the reference lines with which the bootstrap distributions were compared. The pink shaded region, bounded by pink dash-dotted lines, shows the central 95% interval of the reference RoR distribution obtained from the known module-scale distributions of the benchmark generator. The magenta dashed line marks the corresponding theoretical RoR value. The colored annotations above each violin in **b**,**c** report descriptive one-sided empirical bootstrap tail masses ($p$-values) relative to the reference line of the matching color. If the bootstrap median lies below a reference, the displayed value is the fraction of bootstrap samples at or above that reference; if the median lies above it, the lower tail is reported (adaptive). Dashed black brackets below the violins show paired-bootstrap comparisons between module-level estimators (KDE vs GMM) for a fixed per-cell estimator (SAC or CNN). Their two-sided empirical $p$-values were calculated from the paired bootstrap differences relative to zero. Significance is indicated by $^{*}p < 0.05$, $^{**}p < 0.01$, and $^{***}p < 0.001$. In the SAC branch, KDE- and GMM-derived RoR estimates differed significantly in both benchmarks, with GMM yielding larger RoR estimates than KDE (two-sided paired-bootstrap tests).

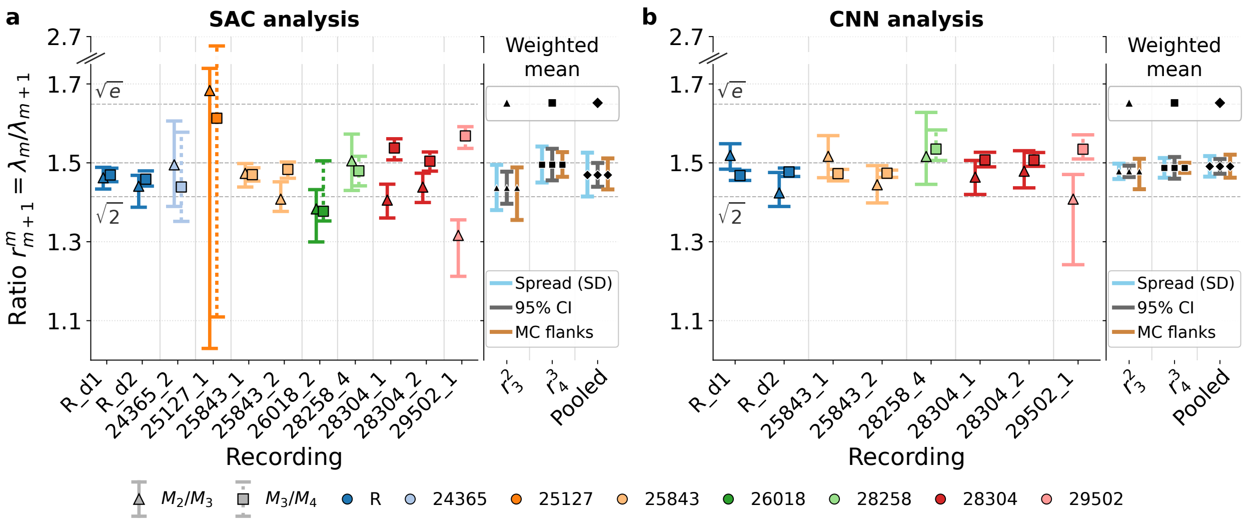


**Supplementary Figure 4. Recording- and population-level adjacent grid-scale ratios.** The two adjacent ratios, $r^2_3$ (triangles) and $r^3_4$ (squares), are plotted for each recording, with central bootstrap 95% intervals, for (**a**) the spatial-autocorrelogram (SAC) analysis and (**b**) the convolutional neural network (CNN) analysis. Symbols mark the raw point estimates obtained from Gaussian mixture models fitted in log space. Recordings are colored by animal; rat R denotes the animal reported by Gardner et al.,[12] and all other animals are from Vollan et al.[13] Dashed horizontal lines mark the $\sqrt{2}$, $3/2$, and $\sqrt{e}$ reference ratios.[9,10]

The analysis includes 11 recordings from 8 animals for the SAC analysis and 8 recordings from 5 animals for the CNN analysis. The population summaries at the right show separate columns for $r^2_3$ and $r^3_4$ and a pooled column containing both ratio types. Repeated recordings from the same animal were first combined so that an animal with several recordings did not contribute several independent population units. All averaging was performed on natural-log ratios and back-transformed for display; the plotted center is therefore a weighted geometric mean.

Recording-level uncertainty was propagated using Monte Carlo sampling. In each of 30,000 repetitions, one value was sampled independently from the bootstrap distribution of every recording belonging to an animal, and the sampled values were combined using fixed reliability weights. Recordings with narrower bootstrap uncertainty received greater weight, with the median uncertainty used as a floor so that an unusually narrow interval could not dominate the animal estimate. Animal MC widths determined the animal reliability weights; the population center and HC1 interval were calculated from the weighted raw animal estimates.

For the pooled column, recordings were first combined separately for $r^2_3$ and $r^3_4$ within each animal. The resulting animal-by-ratio estimates were then entered as separate population units under an independence approximation. This gave $k = 16$ population units for the SAC analysis and $k = 10$ for the CNN analysis. The pooled weighted geometric mean was 1.469 for the SAC analysis, with a descriptive HC1 95% interval of 1.439–1.500, and 1.491 for the CNN analysis (1.472–1.510).

Black symbols mark the weighted population means. The sky-blue "Spread (SD)" intervals show the weighted observed variation among the population units, treated as independent contributions. The gray "95% CI" intervals show the uncertainty in the estimated weighted population mean, calculated using an HC1 heteroskedasticity-consistent standard error and a $t$ distribution with $k - 1$ degrees of freedom. The orange-brown "MC flanks" show the unweighted median lower and upper uncertainty inherited by an animal estimate from the recording-level bootstrap distributions. Thus, the three displays quantify different aspects of uncertainty: variation among population units, precision of the estimated population mean, and the typical recording-level bootstrap uncertainty propagated into an animal estimate.

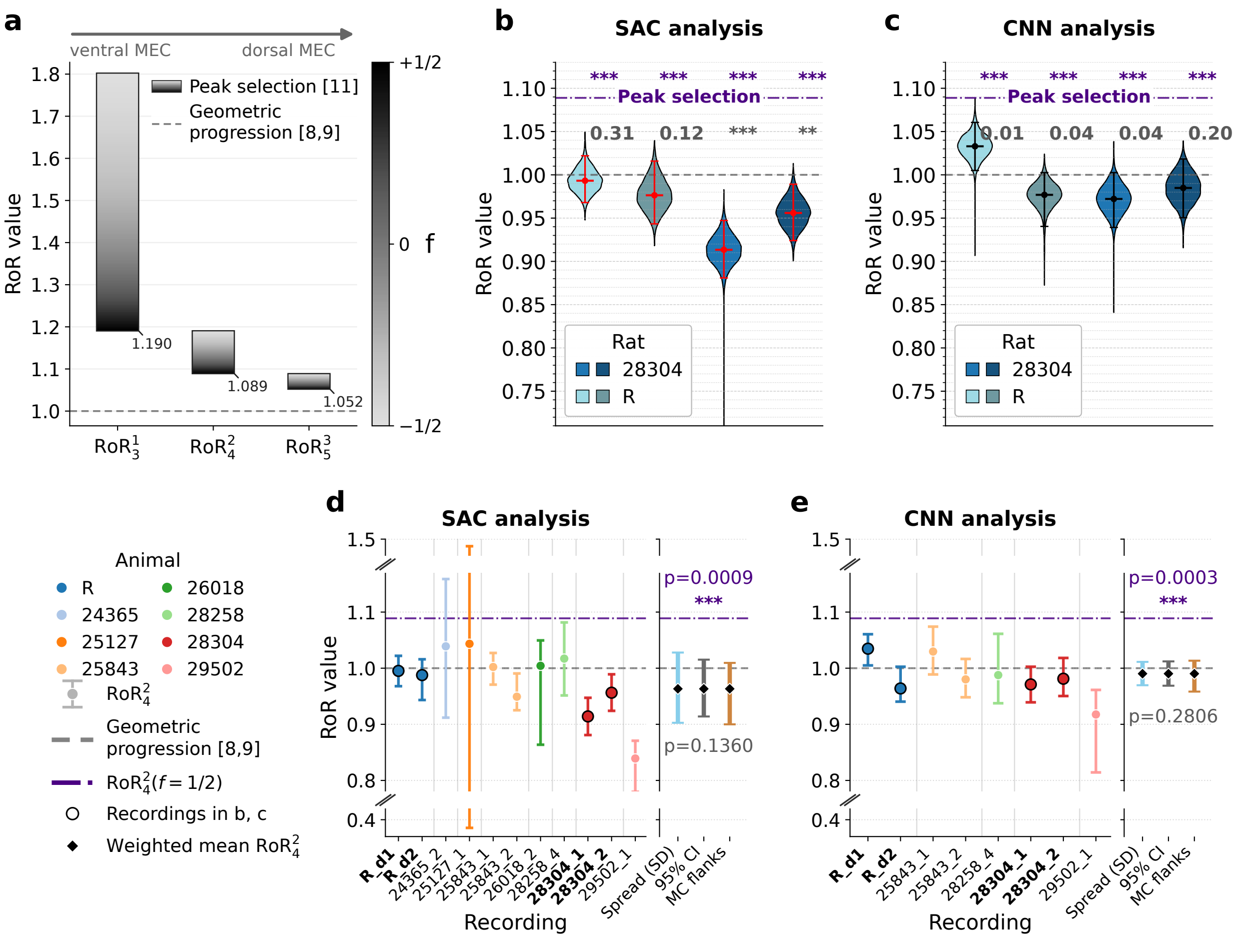


**Supplementary Figure 5. Measured ratio-of-ratios (RoR) values fall below the tested peak-selection bound and lie near unity.** (**a**) Predicted RoR values $\mathrm{RoR}_3^1$, $\mathrm{RoR}_4^2$, and $\mathrm{RoR}_5^3$ for the three consecutive grid-module triples. Geometric-progression theories[8–10] predict RoR = 1 for every module triple (dashed line). Peak selection[11] predicts RoR > 1 for all values of the tunable parameter $f$. For each triple, the prediction is smallest, and therefore most conservative, at $f = \frac{1}{2}$; these lower bounds are $\mathrm{RoR}_3^1 \approx 1.190$, $\mathrm{RoR}_4^2 \approx 1.089$, and $\mathrm{RoR}_5^3 \approx 1.052$. (**b**,**c**) Bootstrap distributions of the measured $\mathrm{RoR}_4^2$ values for four example recordings (rats 25843 and 28304, two recording days each), with grid scale estimated by (**b**) the SAC analysis and (**c**) the CNN analysis. Violins show the retained bootstrap distributions; the marker and whiskers show the bootstrap median and central 95% interval. The gray dashed line marks RoR = 1, and the purple dash-dotted line marks the conservative peak-selection bound $\mathrm{RoR}_4^2(f = \frac{1}{2}) = 49/45 \approx 1.089$.

The colored annotations in **b**,**c** are descriptive one-sided empirical bootstrap tail masses relative to the reference line of matching color. When the bootstrap median lies below a reference, the displayed value is the fraction of retained bootstrap samples at or above that reference; if the median lies above it, the lower tail is reported (adaptive). Because all example medians lie below the peak-selection bound, the purple quantity is $P_{\mathrm{boot}}(\mathrm{RoR}_4^2 \geq 49/45)$. These recording-level bootstrap quantities describe overlap with the reference; they are not the population tests described below. All four example-recording central 95% bootstrap intervals lie below the peak-selection bound. Relative to unity, three of four intervals lie below unity in (**b**; SAC) and one contains unity, whereas in (**c**; CNN) all four intervals contain unity. (**d**,**e**) $\mathrm{RoR}_4^2$ for every recording containing three detected modules, for (**d**) the SAC analysis (11 recordings, 8 animals) and (**e**) the CNN analysis (8 recordings, 5 animals). Rat R was reported by Gardner et al.,[12] all other animals were reported by Vollan et al.[13] Each point is the raw recording estimate with its central bootstrap 95% interval, colored by animal; black outlines identify the recordings shown in (**b**,**c**). Reference lines are as in (**b**,**c**). All 8 intervals in **e** (CNN) and 9 of 11 intervals in **d** (SAC) lie wholly below the conservative peak-selection bound; the broad SAC-analysis intervals for recordings `24365_2` and `25127_1` overlap with that bound (these are the two recordings with the fewest cells – see Supplementary Fig. 1).

The population summaries use the same animal-based procedure as in Fig. 1. Repeated recordings from one animal were first combined, so the final population comparison was made across animals rather than across individual recordings. In each of 30,000 MC repetitions, one RoR value was sampled independently from every recording's bootstrap distribution and the sampled values were combined using fixed reliability weights. This produced one MC uncertainty distribution for each animal. Recordings with narrower bootstrap uncertainty received more weight, but a median-uncertainty floor prevented an unusually narrow recording from dominating. Animals were then combined using the same principle, with weights derived from their MC uncertainty and a second median-uncertainty floor. All calculations were performed in natural-log RoR space and back-transformed for display.

The black diamonds mark the weighted population mean. The sky-blue "Spread (SD)" shows (weighted) observed variation among animals. The gray "95% CI" shows (weighted) uncertainty in the population mean and was calculated using an HC1 (heteroskedasticity-consistent (robust)) standard error and a $t$ distribution with $k-1$ degrees of freedom. The orange-brown "MC flanks" show the (unweighted) typical (median) lower and upper uncertainty of an animal estimate inherited from its recording bootstraps.

The population centers were close to unity: 0.9633 for the SAC analysis (approximate HC1 95% CI, 0.9140–1.0152) and 0.9902 for the CNN analysis (0.9688–1.0121). Both intervals contain RoR = 1 and lie fully below the conservative peak-selection bound $49/45 \approx 1.089$.

The population $p$ values compare the reliability-weighted center with a fixed reference, rather than testing individual recordings. They are approximate two-sided $t$ comparisons in log-RoR space using the HC1 standard error and $k-1$ degrees of freedom. Against RoR = 1, $p = 0.136$ for SAC and $p = 0.281$ for CNN. Against 49/45, $p = 8.91 \times 10^{-4}$ for SAC and $p = 2.72 \times 10^{-4}$ for CNN. These population comparisons are distinct from the descriptive recording-level bootstrap-tail annotations in panels **b** and **c**.
